# Interaction of HS1BP3 with cortactin modulates TKS5 localisation and cancer malignancy

**DOI:** 10.1101/2025.07.14.664640

**Authors:** Arja Arnesen Løchen, Kristiane Søreng, Chiara Veroni, Laura Trachsel-Moncho, Nagham Asp, Robin Gaupset, Helene Knævelsrud, Lars Eftang, Anne Simonsen

**Affiliations:** Department of Molecular Medicine, Institute for Basic Medical Sciences, Faculty of Medicine, University of Oslo, Norway; Centre for Cancer Cell Reprogramming, Institute of Clinical Medicine, Faculty of Medicine, University of Oslo, Norway; Department of Molecular Cell Biology, Institute for Cancer Research, Oslo University Hospital, Norway; Department of Microbiology and Infection Control, Akershus University Hospital, Norway; Department of Digestive Surgery, Akershus University Hospital, Norway

**Keywords:** HS1BP3, cortactin, TKS5, endosomes, proliferation, invasion

## Abstract

We have previously shown that HS1BP3 interacts with the SH3 domain of cortactin, a protein that contributes to a malignant phenotype in cancers. Here we demonstrate that high expression of HS1BP3 is predictive of poorer outcomes for gastric adenocarcinoma and triple negative breast carcinoma patients. We mapped the HS1BP3-SH3 interaction site to the third proline rich region (PRR3.1) of HS1BP3 and show that cells expressing an HS1BP3 PRR3.1 mutant failed to rescue the reduced proliferation and matrix degradation observed in HS1BP3 depleted cells. Moreover, the HS1BP3 PRR3.1 mutant was found to modulate the mRNA levels of the invadopodia scaffold protein TKS5 in gastric cancer cells and contribute to buildup of TKS5 inside multivesicular endosomes in both gastric and TNBC cells. Overall, our results highlight the importance of the direct interaction between the HS1BP3 PRR3.1 and the cortactin SH3 domain in cancer development by regulating endosomal trafficking and cytoskeleton arrangements.

## Introduction

The aggressiveness of cancers is associated with their cell proliferation rate and ability to migrate and degrade extracellular matrix to allow metastasis (1). These attributes are tightly regulated by the interplay between complex molecular pathways, including endosomal trafficking (2) and autophagy (3), as well as several factors involved in cytoskeletal remodelling (4).

Endocytosis involves the internalization of extracellular and plasma membrane localised components into endocytic vesicles that are further sorted for either lysosomal degradation, recycling to the plasma membrane or secretion (2). Several proteins involved in endosomal trafficking have been implicated in tumour metastasis, often associated with the regulation of cellular motility and invasion through the basement layer (2). Endocytosis also crosslinks with the processes of autophagy that involves uptake of cytoplasmic material into double-membraned autophagosome vesicles that fuse with the lysosome for content degradation (5).

We have previously identified and characterised the HCLS1-binding protein 3 (HS1BP3) as a negative regulator of starvation-induced bulk autophagy and hypoxia-induced autophagy of mitochondria (6–8). HS1BP3 localises on recycling endosomes and was found to interact directly with the Src homology 3 domain (SH3) domain of the Src Substrate cortactin (cortactin) (7). Cortactin regulates the actin cytoskeleton through interaction with the actin-related protein 2/3 complex (Arp2/3) and filamentous(F)-actin (9,10) and contains an SH3 domain that interacts with several proteins involved in actin cytoskeleton reorganization (11).

Cortactin is important for cancer cell metastasis through its regulation of actin-cytoskeleton dynamics, permitting cancer cell migration, invadopodia formation and matrix degradation (11–14). Additionally, cortactin regulates endosome trafficking (15,16), the secretion of multivesicular endosome-derived exosomes important for migration (17) and extracellular matrix invasion (18). Cortactin also interacts with the Src substrate and invadopodia scaffold protein TKS5 at the initial stages of invadopodia/podosome formation (14,19).

HS1BP3 has recently been associated with regulation of hepatocellular carcinoma progression (20). We wanted to further examine the role of HS1BP3 in tumorigenesis and investigate if it regulates cancer development via its interaction to cortactin and or by regulation of autophagy or the endocytic pathway. Here we demonstrate that high expression of HS1BP3 may be predictive of poor prognosis in gastric adenocarcinoma and triple negative breast cancer (TNBC). We identify a proline-rich region (PRR) in HS1BP3 important for its direct interaction with Cortactin and show that cancer cell proliferation and matrix substrate invasion, as well as the localization and stability of TKS5, depends on this interaction.

## Results

### High expression of *HS1BP3* is associated with reduced survival of gastric adenocarcinoma and TNBC patients

To investigate a possible role of HS1BP3 in tumorigenesis, we first used the Kaplan-Meier plot analysis tool to address whether *HS1BP3* gene expression levels may predict the survival of cancer patients (21,28). Interestingly, high *HS1BP3* expression was a strong predictor of poor survival of gastric cancer patients (Fig 1A, 1B: hazard ratio HR= 1.64) and patients with the TNBC subtype (Fig 1A, Fig EV1A-B; HR=1.92), an aggressive subtype that is depleted of oestrogen and progesterone receptors and HER2 growth factor (29). *HS1BP3* expression in tumour tissue was also significantly but weakly predictive of improved survival in all subtypes of breast (HR = 0.76) and ovarian (HR = 0.82) cancer patients (Fig 1A, Fig EV1A,C). There were no significant associations between *HS1BP3* expression levels and cancer patient survival in other cancer types. This suggests that HS1BP3 may act as a tumour-promoter in TNBC and gastric cancer specifically.

**Figure 1:**
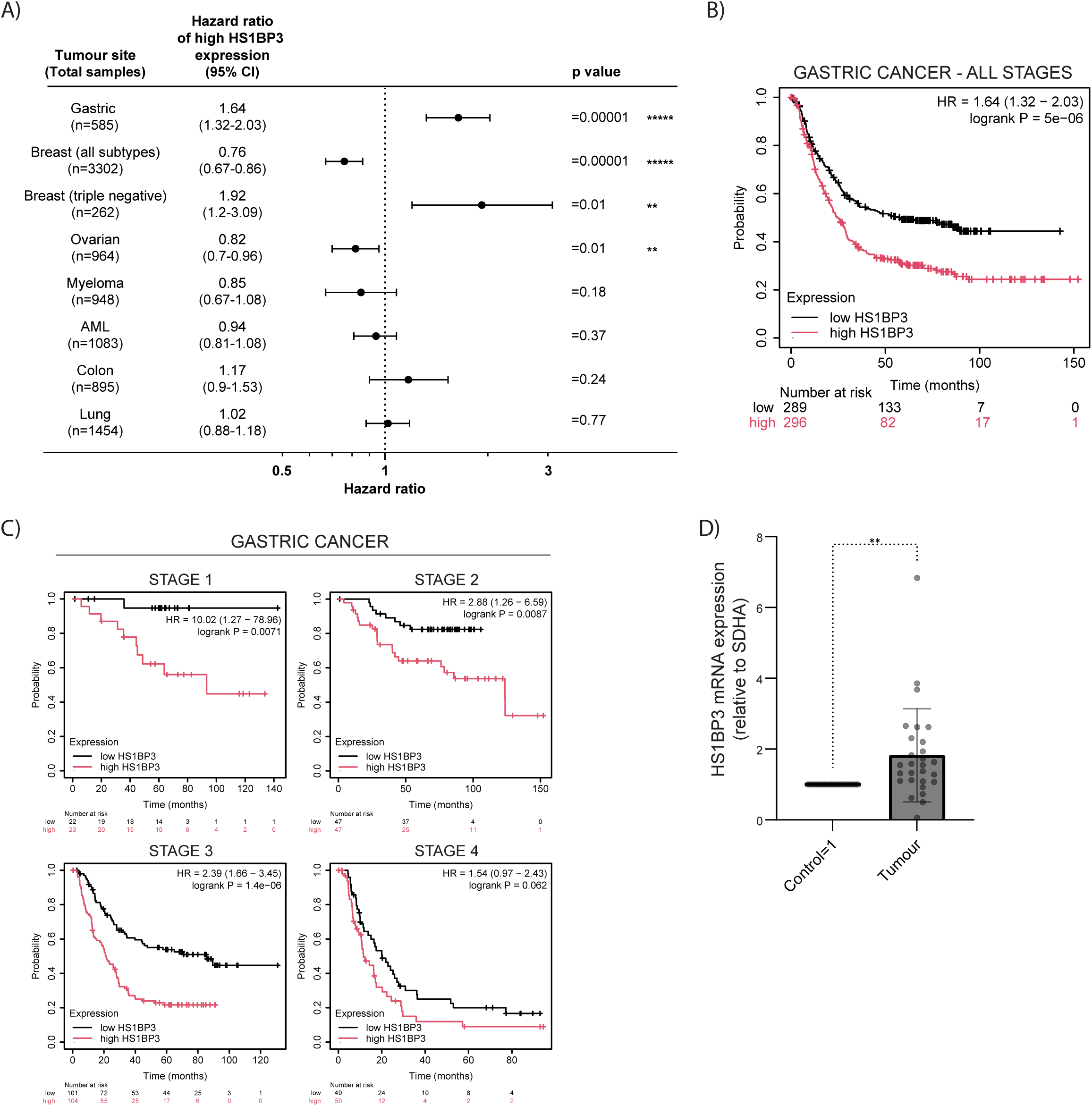
HS1BP3 levels negatively correlates with survival of gastric cancer patients. (**A**) Forest plot showing the hazard ratio plotted on a Log_10_ scale with 95% confidence interval (CI) of high (T3 = third highest expressing tertile) mRNA expression of *HS1BP3* compared to low (T1 = third lowest expressing tertile) *HS1BP3* expressing patients in eight cancer types. The hazard ratio and p-values were calculated using the Cox proportional hazards regression tool in the Kaplan Meier (KM) plotter database. (**B,C**) KM plots showing the probability of survival in patients with gastric cancer according to high (T3 = third highest expressing tertile) and low (T1 = third lowest expressing tertile) mRNA expression of *HS1BP3* in tumour at time of detection at all cancer stages (**B**) and according to cancer stage (**C**). (**D**) Healthy control and tumour gastric tissue from 28 gastric adenocarcinoma patients were analysed for *HS1BP3* mRNA expression using quantitative PCR. P values were determined using one-way ANOVA with Dunnett’s multiple comparison test. **= P <0.01; ***= P <0.001; n.s.= not significant.

When analysing the correlation between *HS1BP3* levels and patient survival with respect to gastric cancer development stage, we found that a high HS1BP3 expression level was a significant and strong predictor of poor outcomes among gastric cancer patients at stage 1-3 (HR= 10.02, 2.88, 2.39 respectively) but not at stage 4 (HR= 1.57, p>0.05), indicating that *HS1BP3* has an important role in the early phase of cancer development (Fig 1C). Using the GTex portal to analyse *HS1BP3* mRNA expression levels in different tissues, we found that *HS1BP3* is highly expressed at the gastro-oesophageal junction and also in stomach tissues (Fig EV1D). Moreover, comparison with the TCGA database, demonstrated that *HS1BP3* is significantly higher expressed in stomach adenocarcinoma and breast carcinoma compared to healthy tissue (Fig EV1E-F).

We next collected gastric tissue from 28 gastric adenocarcinoma patients undergoing gastrectomy surgery, and harvested mRNA from healthy control tissue and the cancer, followed by pair-wise comparison of the mRNA levels of HS1BP3 in each patient (Fig 1D). Overall, these data showed a small but significant increase in *HS1BP3* expression levels in the tumour compared to the healthy control tissue, demonstrating that *HS1BP3* is specifically upregulated in gastric adenocarcinoma.

### HS1BP3 is important for the proliferation of gastric adenocarcinoma cells

To examine the role of HS1BP3 in the early stages of gastric cancer, we acquired three immortalised gastric carcinoma cell lines: AGS, NCI-N87 and MKN-74. While the AGS cell line is derived from a primary gastric adenocarcinoma site, the NCI-N87 and MKN-74 cell lines are sampled from liver metastases of gastric adenocarcinomal origin.

The protein level of HS1BP3 was significantly highest in MKN-74 cells, which also had the highest level of TKS5, while cortactin levels did not differ among the three cell lines (Fig 2A). Intriguingly, a correlation between HS1BP3 and TKS5 expression was also found when studying co-expression among the genes in TCGA and GTEx datasets and in Kaplan-Meier plots. mRNA expression of *HS1BP3* showed a significant and strong positive correlation with the expression of *SH3PXD2A*, the gene encoding TKS5 (Fig EV2A), while HS1BP3 negatively correlated with *CTTN*, the gene encoding cortactin (Fig EV2B). In Kaplan-Meier plots, high *SH3PXD2A* levels predicted poorer outcomes of gastric cancer patients but not for TNBC patients (Fig EV2C-D), while *CTTN* expression levels did not predict survival outcomes of neither gastric nor TNBC patients (Fig EV2E-F). Together this indicates a positive correlation between HS1BP3 and TKS5 expression.

**Figure 2:**
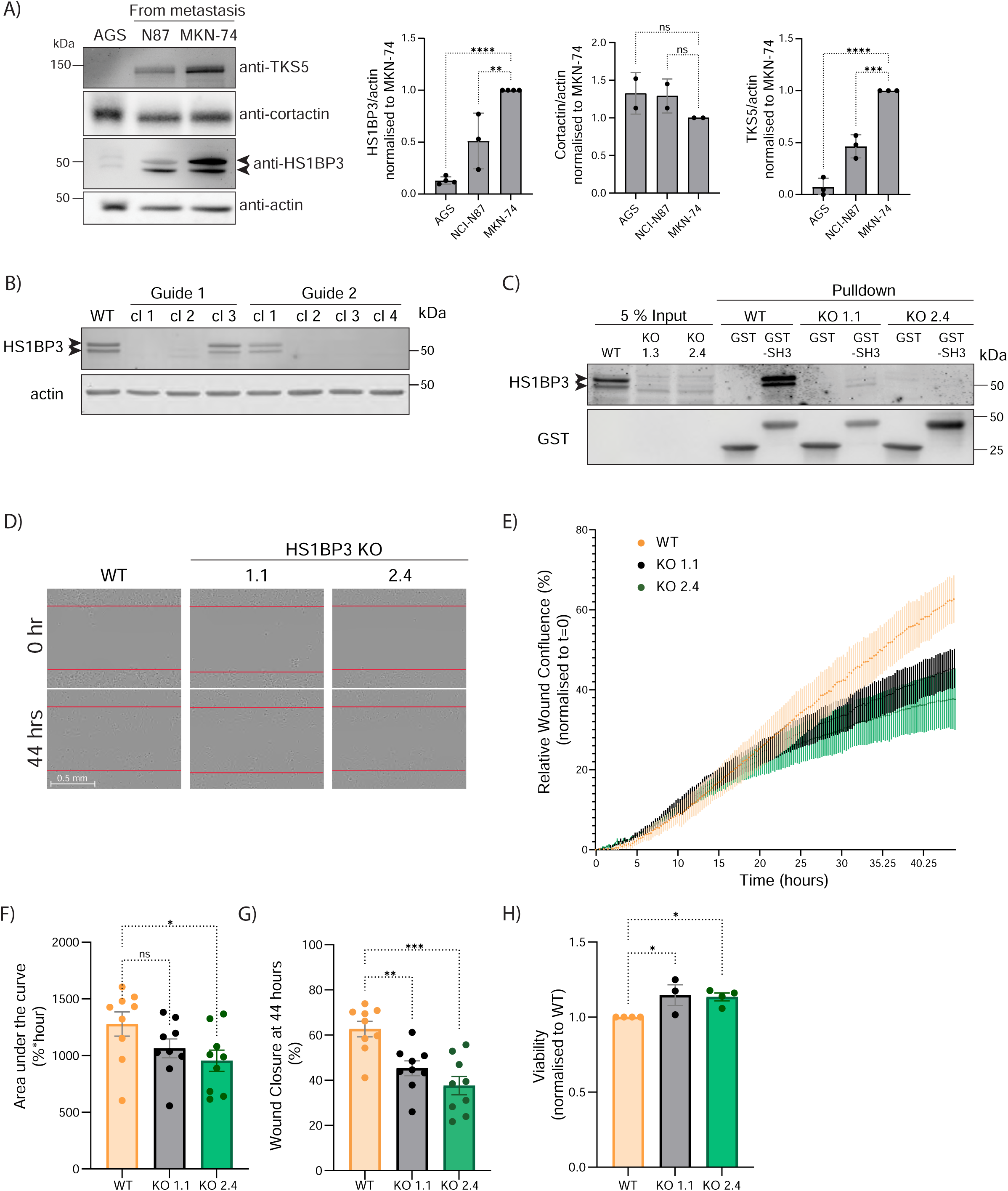
HS1BP3 promotes migration of a gastric cancer cell line. (**A**) Immunoblotting and quantification of protein levels of TKS5, cortactin, and HS1BP3 in three gastric cancer cell lines (AGS, N87 and MKN-74). Quantification shows the mean level of the indicated protein, and error bars indicate the standard deviation of the mean. N=2-4. (**B**) MKN-74 wild type (WT) cells and single MKN-74 clones subjected to CRISPR/Cas9 knockout (KO) of HS1BP3 using two different gRNA (guide 1 and guide 2) were subjected to immunoblotting against HS1BP3. Actin was used as a loading control, n=1. (**C**) GST-SH3 (SH3 domain of cortactin) pulldown of HS1BP3 from MKN-74 lysates of WT and two single HS1BP3 KO clones (clone 1.1 and 2.4). n=1. (**D-E**) MKN-74 WT and HS1BP3 KO clones (1.1 and 2.4) were seeded confluently in 96-well plates and left overnight before creating wounds and imaged by live cell microscopy every ten minutes over 44 hours to quantify the filling of the wound. (**D**) Representative images of cells at 0 hours or 44 hours after wound scratch (**E**) Plots show the relative wound confluence of each well up to 44 hours after wound scratch, n=3, (**F**) the area under the curve of (E) and (**G**) final wound confluence at 44 hours of (E). (**H**) Cell viability of MKN-74 WT and HS1BP3 KO clones (1.1. and 2.4) cells, measured by the MTT metabolic assay. The data shown are mean ± standard error of the mean. All P values were determined using one-way ANOVA with Dunnett’s multiple comparison test. *= P <0.05; **= P <0.01; ***= P <0.001; ; ****= P<0.0001; n.s.= not significant.

To further understand the role of HS1BP3 in tumorigenesis, we generated HS1BP3 knockout (KO) MKN-74 cells using two different CRISPR/Cas9 guide RNAs targeting either exon 1 or 2 (guide 1 and guide 2, respectively) of the HS1BP3 gene. One cell clone per guide was selected for further experiments, both showing efficient depletion of HS1BP3 (guide 1 clone 1=1.1; and guide 2 clone 4=2.4) (Fig 2B). GST pulldown experiments with the SH3 domain of cortactin, showed strong binding of HS1BP3 using cell lysates from MKN-74 wild-type (WT) cells, while no HS1BP3 was detected in cell lysates from KO clone 2.4 and only a small amount of HS1BP3 was present in KO clone 1.1, demonstrating that HS1BP3 was completely abolished in clone 2.4 (Fig 2C). Using a wound-healing assay to study directed cell migration, we observed a significantly reduced level of wound closure in both HS1BP3 KO clones compared to MKN-74 WT cells (Fig 2D-G). Clone 1.1 had a slightly lower reduction in migration compared to clone 2.4, which may suggest that a small amount of HS1BP3 is enough to drive the HS1BP3-mediated cell migration. It should be noted that the MKN-74 cells migrate slowly, as the wound did not close completely within 50 hours of culturing, and a difference in migration between WT and KO cells was only observed after 20 hours. Hence, it is likely that a difference in proliferation, rather than migration, explains the observed difference between MKN-74 WT and HS1BP3 KO cells in the wound assay. To our surprise, both HS1BP3 KO clones were significantly associated with approximately 15% higher metabolic activity or viability compared to WT cells, as analysed in an MTT assay (Fig 2H), indicating that the reduced proliferation or migration of HS1BP3 KO cells were not due to increased cell death.

### HS1BP3 directly interacts with cortactin via its third proline-rich region PRR3.1

Previous publications report that stable overexpression of cortactin in cancer cell lines is associated with an invasive phenotype (9,10). Although *CTTN* expression levels did not predict survival outcomes of gastric cancer patients (Fig EV2E), the interaction of HS1BP3 with cortactin may be important for HS1BP3’s role in gastric cancer development. We previously demonstrated that HS1BP3 interacts with the SH3 domain of cortactin (7) but did not determine the HS1BP3 region responsible for its interaction with cortactin. PRRs commonly interact with SH3 domains (11,30), and we speculated that HS1BP3 binds to cortactin’s SH3 domain via one or more of its PRRs. To address this, we first generated HS1BP3-SH3 docking models using the AlphaFold Server (Fig 3A). The five resulting models suggested that HS1BP3 interacts with the cortactin SH3 domain via its lipid-binding PX domain and a region spanning its large C-terminal PRR3 (Fig 3A-C). We have previously shown that the PX domain is indispensable for the HS1BP3-cortactin interaction (7), suggesting limitations of the AlphaFold model. We therefore decided to generate and purify four MBP-tagged HS1BP3 C-terminal deletion mutants (Fig 3B) that were incubated with the SH3 domain of cortactin fused to GST and bound to glutathione beads (Fig 3D-E). While the full-length MBP-HS1BP3 bound strongly to GST-SH3, the deletion mutants HS1BP3^PRR1^ and HS1BP3^PRR1-2^ containing the PX domain and PRR1 or PRR1-2, respectively, did not interact with GST-SH3 (Fig 3D-E). Since the AlphaFold models indicated that the SH3 domain interaction with the HS1BP3 PRR3 region may vary between two potential PRRs (PRR3.1 and PRR3.2) (Fig 3A), we decided to make C-terminal deletion mutants containing either PRR3.1 (HS1BP3^PRR1-3.1^) or PRR3.2 (HS1BP3^PRR1-3.2^) (Fig 3B). Both these deletion mutants bound to GST-SH3 but the interaction with HS1BP3^PRR1-3.1^, containing PRR1-3.1, was strongly reduced (Fig 3D-E). Overall, our data indicate that the PX domain, PRR1 and PRR2 are dispensable for the direct interaction between HS1BP3 and the SH3 domain of cortactin, while PRR3.1 and PRR3.2 are the likely interaction regions.

**Figure 3:**
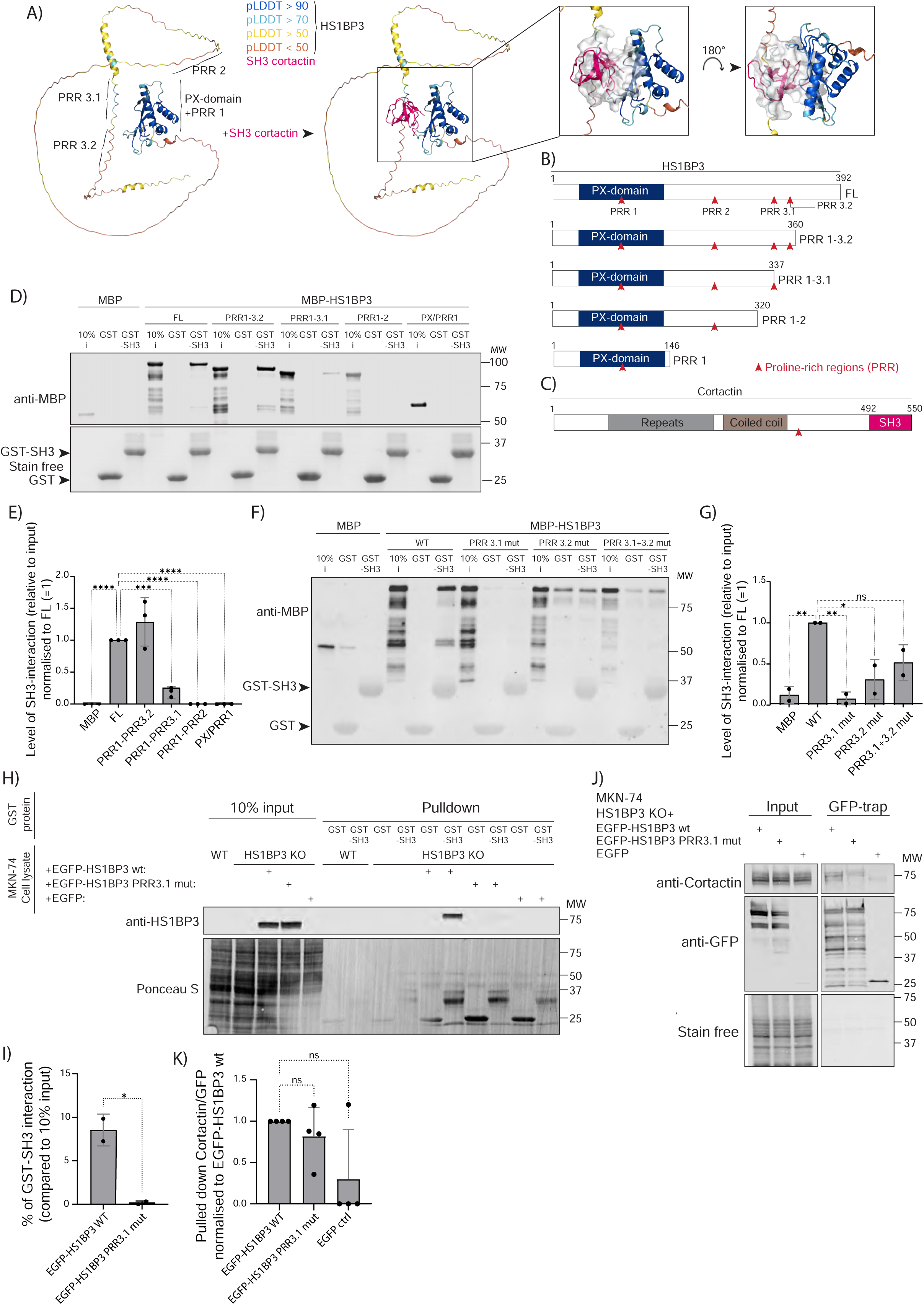
HS1BP3 interacts directly with the SH3-domain of cortactin via its third proline-rich region PRR3.1. (**A**) An AlphaFold-generated prediction on the interaction between HS1BP3 and the SH3 domain of cortactin. The per-residue measures of local confidence (Plddt) values are scaled 0-100 and indicate confidence of the predictions. The cloud indicates interaction surfaces. (**B**) The cartoon shows the Full length (FL) HS1BP3 and position of the HS1BP3 PX domain and the proline-rich regions (PRR). The C-terminal deletion mutants used to map the interaction of HS1BP3 with the SH3 domain of cortactin are shown. (**C**) The cartoon indicates the F-actin binding repeats region, the coiled coil, PRR and SH3 domain of cortactin. (**D**) GST-SH3 pulldown assay using recombinant GST, GST-SH3 from cortactin, and MBP-HS1BP3 full length and deletion mutants, followed by immunoblot analysis of bound HS1BP3 proteins using an anti-MBP antibody. GST and GST-SH3 were visualized using stain-free gels. 10% of the input (i) MBP protein was loaded. n=3. (**E**) Quantification of the data in (D), showing the mean level of HS1BP3 interaction to GST-SH3. (**F**) GST-pulldown assay of recombinant MBP-HS1BP3 full-length WT and mutants of PRR3.1 and/or PRR3.2, followed by immunoblot analysis of bound HS1BP3 proteins using an anti-MBP antibody. The levels of GST and GST-SH3 used are seen. 10% of the input (i) MBP protein was loaded. n=2. (**G**) Quantification of the data in (F) showing the mean level of HS1BP3 interaction to GST-SH3. (**H**) GST-pulldown assay of cell lysates from MKN-74 WT cells or HS1BP3 KO cells rescued or not with EGFP-HS1BP3 WT, EGFP-HS1BP3 PRR3.1-mutant or EGFP-control, followed by anti-HS1BP3 immunoblot analysis to determine binding to GST and GST-SH3 (SH3 domain of cortactin), n=2. (**I**) Quantification of the data in (H), showing the mean percentage EGFP-HS1BP3 binding to GST-SH3, n=2. The levels of GST and GST-SH3 were detected by Ponceau S staining of the membrane. (**J**) Cell lysates from MKN-74 cells expressing EGFP-HS1BP3 WT, PRR3.1-mutant or EGFP-control were subjected to GFP-trap pulldown and immunoblot analysis of cortactin, n=4 (**K**) quantification of (J) showing the mean percentage pulldown of cortactin relative to pulled down GFP. Data are mean ± standard error of the mean. P-values were determined using one-way ANOVA with Dunnett’s multiple comparison test. *= P <0.05; **= P <0.01; ***= P <0.001; ****= P<0.0001; n.s.= not significant.

To further validate a role for HS1BP3 PPR3 in binding to the SH3 domain of cortactin, we produced full-length MBP-HS1BP3 having the prolines of PRR3.1 and PRR3.2 replaced with alanine, either separately or together (Fig 3F-G). Intriguingly, mutation of PRR3.1 completely prevented the binding between MBP-HS1BP3 and GST-SH3. Mutation of PRR3.2 induced non-specific interactions of both the single and double mutant, indicating it may induce HS1BP3 conformational changes. Since the HS1BP3 PRR3.1-mutation prevented the direct interaction between MBP-HS1BP3 and GST-SH3 *in vitro*, we next set out to investigate the role of HS1BP3-PRR3.1 in cells. The MKN-74 HS1BP3 KO cells were rescued with EGFP-tagged WT or PRR3.1-mutant HS1BP3, followed by incubation of cell lysates with GST or GST-SH3 (cortactin). Indeed, immunoblot analysis of bound HS1BP3 revealed that binding of the HS1BP3 PRR3.1 mutant to GST-SH3 was strongly reduced compared to HS1BP3 WT (Fig 3H-I). Co-immunoprecipitation of endogenous cortactin with EGFP-HS1BP3 WT or PRR3.1 mutant demonstrated that the PRR3.1 mutant can still interact with cortactin in cells, but at a reduced level (Fig 3J-K). We postulate that this may be due to indirect interactions of HS1BP3 with cortactin in a protein complex, since there are at least 221 SH3-domain containing proteins that can link them together (30).

Taken together, our data indicate that the PRR3 region of HS1BP3 facilitates its interaction with the SH3 domain of cortactin, where mutation of PRR3.1 in full-length HS1BP3 prevents its direct interaction with cortactin’s SH3 domain.

### The direct HS1BP3-cortactin interaction modulates cancer cell proliferation and invasion

As the EGFP-tagged HS1BP3 WT and PRR3.1-mutant proteins were expressed at a much higher level than the endogenous HS1BP3 protein (Fig 3H), we decided to generate new MKN-74 HS1BP3 KO cells stably expressing untagged HS1BP3 WT and PRR3.1-mutant at a level close to the endogenous protein (Fig 4A). These cells, together with MKN-74 WT and HS1BP3 KO cells were then used to determine if the HS1BP3-cortactin interaction was important for cell proliferation. In line with the wound-healing assay (Fig 2D-G), we found that the proliferation of HS1BP3 KO cells was reduced compared to WT cells when quantifying cell confluence over 51 hours (Fig 4B-D). Intriguingly, HS1BP3 WT but not the PRR3.1 mutant was able to rescue the reduced proliferation of HS1BP3 KO cells (Fig 4B-D), indicating that the interaction of HS1BP3 with cortactin promotes cell proliferation.

**Figure 4:**
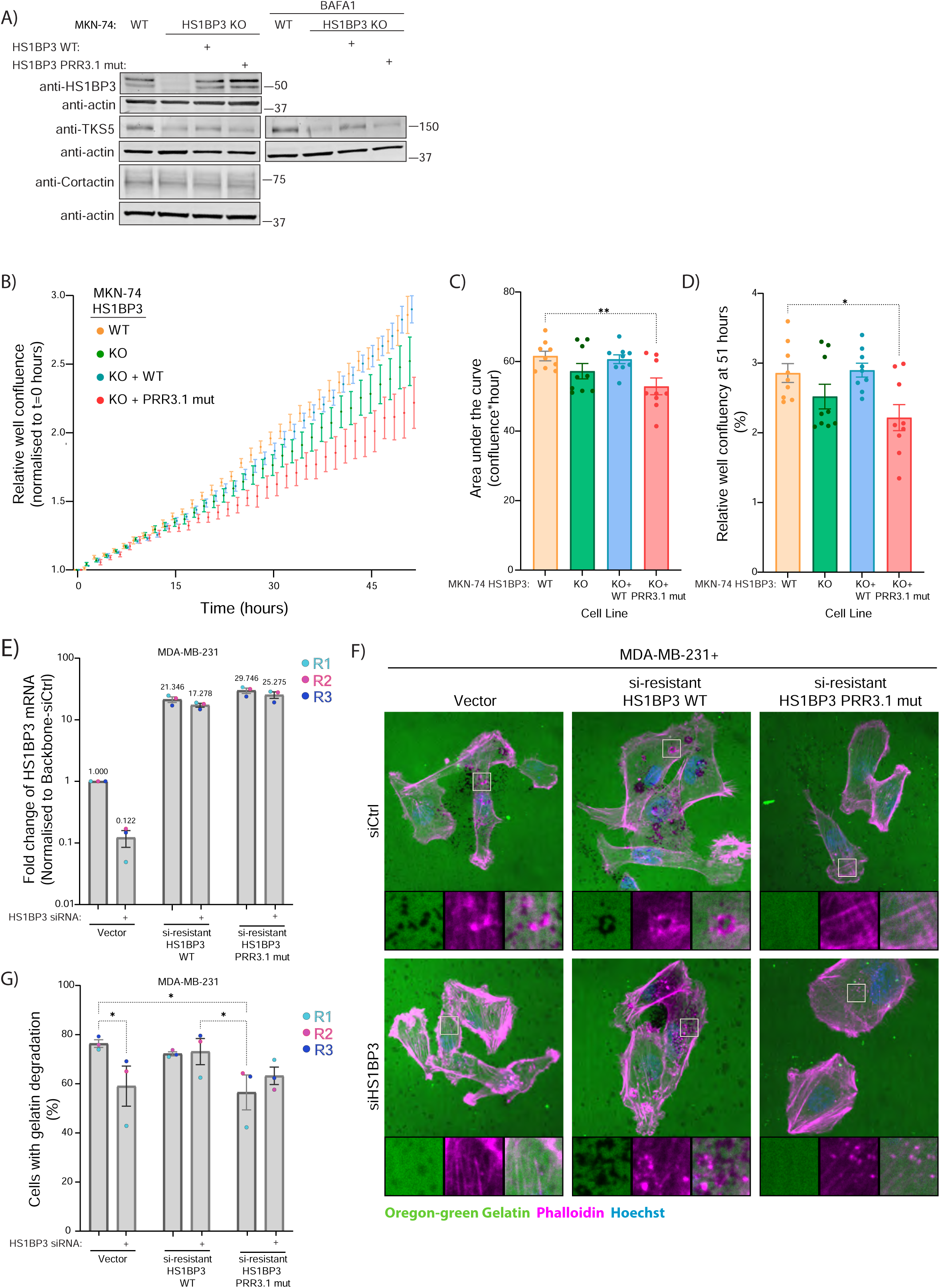
The HS1BP3-cortactin interaction modulates gastric cell proliferation and invasion in TNBC. (**A**) Cell lysates from MKN-74 WT, HS1BP3 KO or HS1BP3 KO with rescue with low-expressing HS1BP3 WT or PRR3.1-mutant were treated or not for 2 hours with 100 nM Bafilomycin A1 (BafA1) followed by immunoblot analysis for cortactin and TKS5 levels, n=3. (**B**) MKN-74 WT, HS1BP3 KO 2.4 and KO 2.4 with rescue of HS1BP3 WT or PRR.1-mutant were seeded sparsely in 96-well plates for live cell microscopy to observe their proliferation rate as mean relative confluence every hour over 51 hours, n=3. (**C-D**) Quantifications of (B) showing the area under the curve (**C**) and relative well confluency after 51 hours (**D**). Each data point corresponds to a single well (technical replicate). (**E-G**) MDA-MB-231 stably expressing negative-control vector, siRNA-resistant HS1BP3 WT or PRR3.1-mutant were transfected with negative-control siRNA or siRNA targeting endogenous HS1BP3. 72 hours after transfection the cells were harvested and checked for HS1BP3 expression using quantitative PCR (**E**) or seeded sparsely on coverslips coated with Oregon-Green labelled gelatin for 4 hours before PFA-fixation and staining with Hoechst and phalloidin before visualising the cells by confocal imaging (**F**). Images are taken with a Nikon Ti2-E microscope with a Yokogawa CSU-W1 SoRa spinning disk 60xWI water immersion objective. Inset rectangles are 9x9 µm^2^. (**E**) data are plotted on Log _10_ scale for easier visualisation and comparison of values. The mean expression value normalised to Vector siCtrl are written on top of the bar. Each data point corresponds to one replicate (**G**) quantification of (F) showing the percentage of quantified cells with observed gelatin degradation spots within cell boundary compared to total number of quantified cells. Each data point corresponds to one replicate quantifying between 61-165 cells. Data are mean ± standard error of the mean. P-values were determined using one-way ANOVA with Dunnett’s multiple comparison test (C-D) or two-way ANOVA with Holm-Šídák (E) or Fisher’s LSD (G) test. *= P <0.05; **= P<0.01; no significance line = not significant.

Since HS1BP3 has been found to act as a negative regulator of autophagy and autophagy levels can affect cancer development (3,7), we aimed to determine if HS1BP3 affects proliferation of MKN-74 cells via regulation of autophagy. By quantifying the level of autophagosomes (LC3B puncta) and autolysosomes (LC3B-LAMP1 overlap) in immunofluorescence images (Fig EV3A-D) and measuring LC3B flux by western blot (Figure S3E-H) in control and starved conditions, we did, however, not detect any significant differences in autophagy levels in HS1BP3 KO cells versus HS1BP3 WT or PRR3.1 rescue cells. Thus, the interaction of HS1BP3 with cortactin seems not to regulate autophagy in gastric adenocarcinoma cells.

To further assess whether the HS1BP3-cortactin interaction may play a role in cancer cell invasion, MKN-74 cells were seeded on Oregon-Green labelled gelatin for up to 17 hours, followed by imaging analysis. Unfortunately, we did not detect any convincing matrix degradation of the MKN-74 WT cells (Fig EV4A), suggesting that these cells are not suited for invasion assays.

As high *HS1BP3* expression also was found to correlate with poor survival of patients with TNBC (Fig 1A, Fig EV1B; HR=1.92), we decided to use the TNBC cell line MDA-MB-231 to assess the role of HS1BP3 in cancer cell invasion. MDA-MB-231 is a cell line with adenocarcinomal histology that robustly degrades gelatin (31). Intriguingly, siRNA mediated depletion of HS1BP3 in MDA-MB-231 cells (Figure 4E), caused a significant reduction in the number of cells with areas of gelatin degradation after 4 hours of incubation on Oregon-Green Gelatin compared to cells transfected with control siRNA (Fig 4F-G). The gelatin degradation was rescued in cells with stable expression of siRNA-resistant HS1BP3 WT but not in cells expressing siRNA-resistant HS1BP3 PRR3.1-mutant upon knockdown of HS1BP3 (Fig 4F-G). Taken together, our data demonstrate that the interaction of HS1BP3 with cortactin is important for cancer cell proliferation and invasion.

### The HS1BP3-cortactin interaction regulates levels of TKS5-associated multivesicular endosome positioning and secretion of TKS5

To further understand why the HS1BP3-cortactin interaction modulates cancer development, we decided to follow up on our initial finding of a correlation between HS1BP3 and TKS5 expression levels in gastric cancer cell lines (Fig 2A). Intriguingly, we observed a reduced level of TKS5 in the MKN-74 HS1BP3 KO cells, which was rescued by HS1BP3 WT but not the PRR3.1-mutant (Fig 4A). TKS5 did not accumulate upon treatment with the lysosomal V-ATPase inhibitor Bafilomycin A1 (Fig 4A), indicating that cells lacking HS1BP3 or being unable to interact with cortactin may have lower levels of TKS5 expression. Indeed, the mRNA level of *SH3PXD2A* (TKS5) was reduced in MKN-74 HS1BP3 KO cells compared to WT and HS1BP3 WT rescue cells (Fig EV4B). In contrast, HS1BP3 KO and WT rescue did not affect cortactin mRNA levels (Fig EV4C), in line with the expression-correlation patterns described above (Fig 2A; Fig EV2A-B). Overall, our data suggest that the direct interaction of HS1BP3 with cortactin may affect TKS5 protein levels.

To determine if HS1BP3 may regulate the cortactin-TKS5 interaction, MKN-74 WT and HS1BP3 KO cells were transfected with TKS5-EGFP, followed by GFP pulldown and immunoblot analysis for cortactin and HS1BP3, showing that HS1BP3 neither affected the TKS5-cortactin interaction nor interacted with TKS5 (Fig EV4D-E). We then postulated that HS1BP3 may modulate the subcellular localisation of cortactin and/or TKS5. Hence, MKN-74 WT and HS1BP3 KO cells, rescued or not with HS1BP3 WT or the PRR3.1-mutant, were stably transduced with TKS5-GFP and mCherry-cortactin. The cells were analysed by fluorescence microscopy for the colocalisation and/or changes to TKS5 and cortactin puncta (Fig 5). It is well established that invasive cell lines that degrade extracellular matrix form ventral invadopodia when grown on glass coverslips due to cell matrix secretion (17,31). In line with MKN-74 cells not being invasive when grown on gelatin (Fig EV4A), we were unable to detect any TKS5-GFP puncta ventrally. However, we observed a significant increase in the number of TKS5-GFP puncta in the middle of the cell in the HS1BP3 KO MKN-74 cells rescued with the HS1BP3 PRR3.1-mutant compared to WT HS1BP3 (Fig 5A-C). While we did not observe any significant difference in the percentage of cells with mCherry-cortactin puncta or in the number of mCherry-cortactin puncta per cell between the MKN-74 genotypes (Fig 5D-E), there was a trend of increased TKS5-GFP colocalisation with mCherry-cortactin in MKN-74 WT cells and in HS1BP3 KO rescue with HS1BP3 WT (Fig 5F-G).

**Figure 5:**
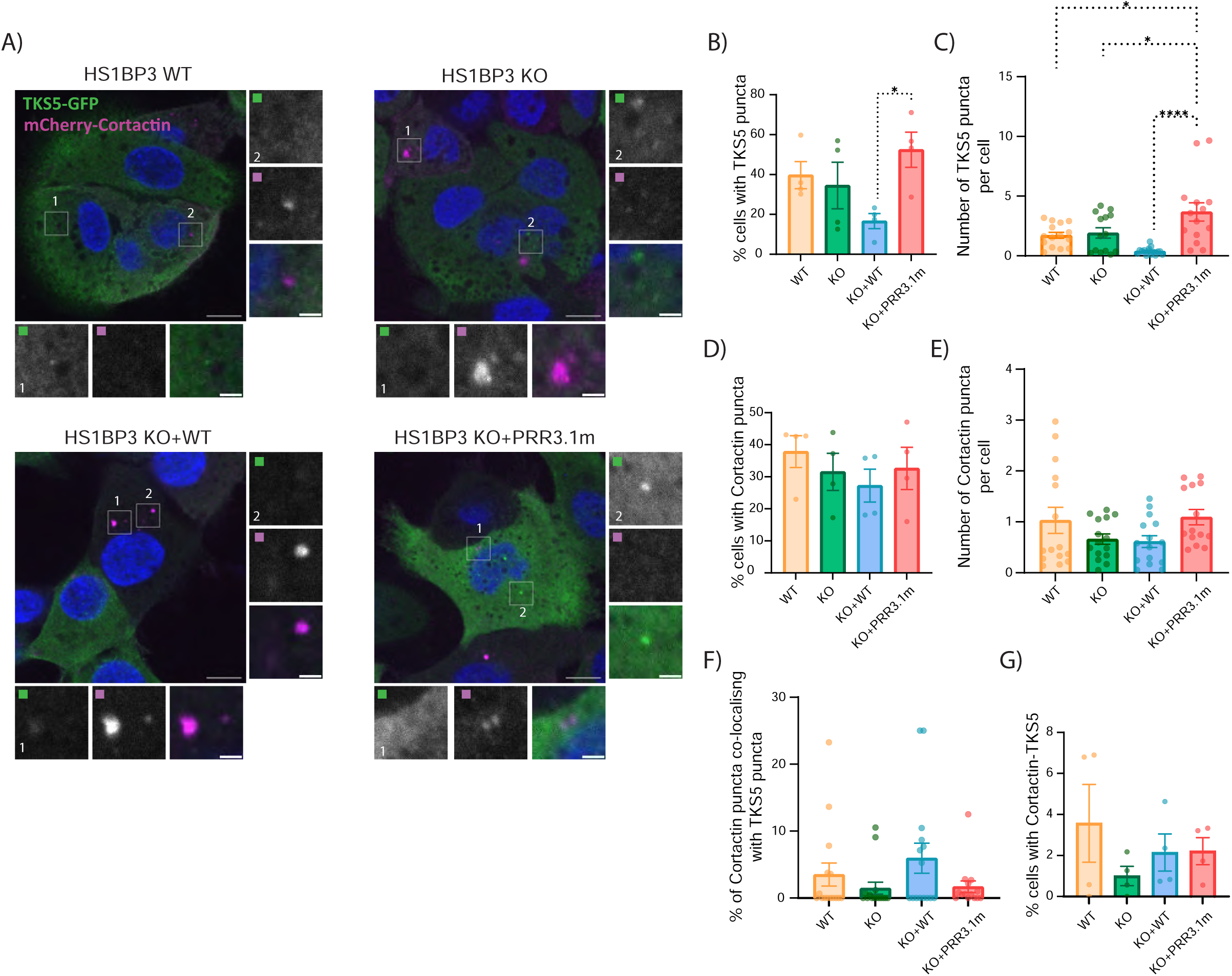
Ablation of the HS1BP3-cortactin interaction is associated with increased levels of TKS5 puncta but does not affect TKS5-cortactin interaction. (**A**) representative fluorescence imaging of overexpressed TKS5-GFP and mCherry-Cortactin in the middle of MKN-74 HS1BP3 WT, KO with or without rescue of HS1BP3 WT or PRR3.1. Images were acquired with a Nikon CREST X-Light V3 spinning disk microscope using a 60× oil objective (NA 1.42). Scale bars: 10 µm (main figure) or 2 µm (inset). (**B-G**) quantification of (A) representing the percentage of cells with at least one puncta of TKS5 (**B**), cortactin (**D**) or cortactin overlapping with TKS5 (**G**), or the number of TKS5 (**C**) or cortactin puncta (**E**) per cell. (**F**) shows the percentage cortactin puncta that overlap with TKS5 puncta. Each data point corresponds to a single field of view. Data are mean ± Standard error of the mean, the statistical significance was calculated with ordinary one-way ANOVA, followed by Tukey’s multiple comparison test (n = 4, 103-238 cells were quantified in each condition). In D-G no statistical significance was observed in the comparisons.

Since we observed an increase in TKS5-GFP puncta in the MKN-74 PRR3.1 rescue cells with few co-localising with cortactin, we postulated that these puncta may represent TKS5 in multivesicular endosomes, from where TKS5 is secreted with metalloproteases important for matrix degradation (25,32). Using MDA-MB-231 cells stably overexpressing TKS5-GFP, we indeed observed a consistent localisation of TKS5-GFP in LAMP1-structures observed as bright puncta (Fig 6A). We did not observe any TKS5-GFP puncta outside LAMP1-positive structures. The fact that the TKS5-GFP signal was not quenched by the low pH of lysosomes, indicates that TKS5-GFP is found within the intraluminal vesicles of LAMP1-positive multivesicular bodies. Intriguingly, while overexpression of HS1BP3 WT significantly decreased the occurrence of TKS5-GFP puncta, overexpression of the PRR3.1 mutant or siRNA-mediated knockdown of HS1BP3 significantly increased the number of TKS5-puncta (Fig 6A-B). The increased number of TKS5-puncta observed in HS1BP3-depleted cells was significantly reduced by overexpression of siRNA-resistant HS1BP3 WT or the PRR3.1 mutant (Fig 6A-B). It is interesting to note that the changes in TKS5-GFP puncta levels occurred without affecting the total TKS5-GFP or endogenous TKS5 levels, as observed by western blot (Fig 6C). Since the number of TKS5-GFP puncta was strongly increased in cells expressing the HS1BP3 PRR3.1 mutant, we speculated that this may be associated with inhibited secretion of TKS5 in multivesicular bodies. To test this, we collected conditioned media from parental and HS1BP3 WT and PRR3.1 overexpressing MDA-MB-231 cells and used 100 K spin-columns to enrich the media of extracellular vesicles. From this we observed a slightly reduced level of secreted TKS5 from PRR3.1 overexpressing cells (Fig 6D).

**Figure 6:**
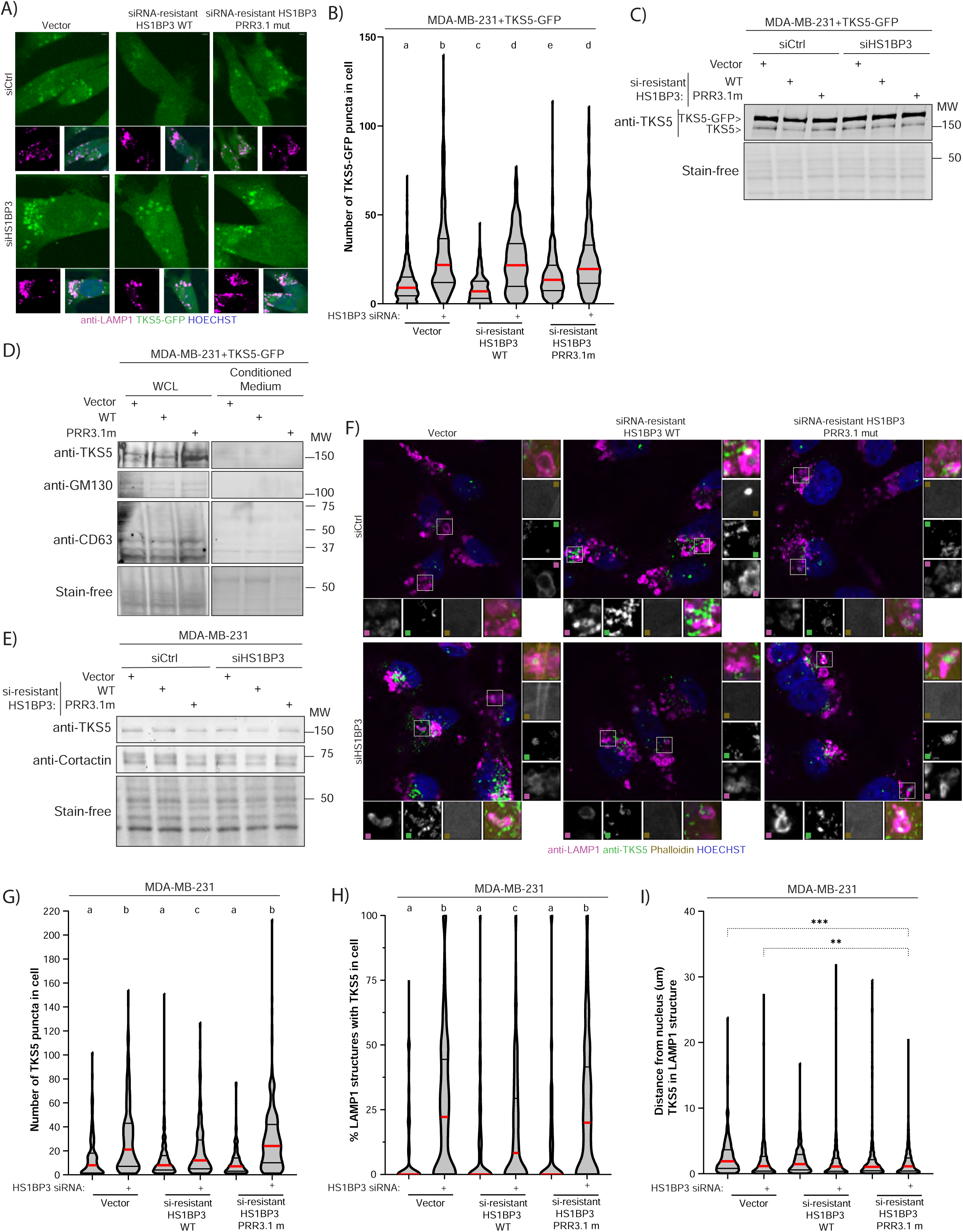
The HS1BP3-cortactin interaction regulates levels of TKS5-associated multivesicular endosome and secretion of TKS5. (**A**) Representative immunofluorescence images of MDA-MB-231 with stable expression of TKS5-GFP and negative control vector or siRNA-resistant HS1BP3 WT or PRR3.1-mutant. The cells were subjected to transient transfection with a negative siRNA-control or siRNA targeting HS1BP3 and were PFA-fixed after 72 hours of knockdown. This was followed by antibody staining with LAMP1 and imaging. Images are taken with a Nikon Ti2-E microscope with a Yokogawa CSU-W1 SoRa spinning disk 60xWI water immersion objective. Scale bar is 2 µm. (**B**) violin plot with quantification of (A) number of segmented TKS5-GFP puncta in all segmented cells. 44-660 cells were quantified in each condition. (**C**) Cells treated as in (A) were lysed instead of PFA-fixed and total TKS5 (TKS5-GFP and endogenous) were analysed by western blot with total protein level (stain-free) as a loading control. Representative of three experiments. (**D**) MDA-MB-231 with stable expression of TKS5-GFP with negative control vector or siRNA-resistant HS1BP3 WT or PRR3.1-mutant were seeded in 150 mm dishes and serum-starved for 44 hours before the conditioned media was collected and extracellular vesicles concentrated using a 100 K spin column. 10% of the whole cell lysate (WCL) and 8% of the concentrated media were analysed by western blot with indicated antibodies and total protein level (stain-free) as loading control. (**F**) Representative immunofluorescence images of MDA-MB-231 with negative control vector or siRNA-resistant HS1BP3 WT or PRR3.1-mutant. The cells were subjected to transient transfection with a negative siRNA-control or siRNA targeting HS1BP3 and were PFA-fixed after 72 hours of knockdown. This was followed by antibody staining with anti-TKS5 and anti-LAMP1, and imaging. Images are taken with a Nikon Ti2-E microscope with a Yokogawa CSU-W1 SoRa spinning disk 60xWI water immersion objective. Inset is 5.5x5.5 µm. (**G**) Cells treated as in (E) were lysed instead of PFA-fixed and total TKS5, cortactin and total protein (stain-free) were analysed by western blot with total protein level (stain-free) as a loading control. (**G, H, I**) violin plots with quantifications of (F) showing the (G) quantified number of TKS5 in cells, (H) the % of LAMP1 structures in cells that contain TKS5, and (I) the distance between nucleus and LAMP1-structures containing TKS5. 47-191 cells were quantified in each condition. In (B, G-I) the thick red line is the median and the black horizontal lines are the quartiles. Significant difference (p<0.05) is indicated on basis of different letter (a-e) between the condition (in conditions with the same letter, the difference between the means is not statistically significant), or **=<0.01 or ***=<0.001. The statistical significance was calculated with ordinary one-way ANOVA followed by Holm-Šídák multiple comparison test.

To test if HS1BP3 expression levels also have similar effects on endogenous TKS5, MDA-MB-231 cells with or without expression of siRNA-resistant HS1BP3 WT or PRR3.1 mutant were depleted or not of HS1BP3, followed by immunoblotting analysis. Similarly to overexpressed TKS5-GFP (Fig 6C), the total levels of endogenous TKS5 did not change between the conditions (Fig 6E). We then stained fixed cells for endogenous TKS5 and LAMP1 to look at recruitment of TKS5 to LAMP1-positive structures (Fig 6F). Although endosomal localization of endogenous TKS5 is a rare event compared to overexpressed TKS5-GFP (33,25), we observed a significant increase of TKS5 puncta per cell in HS1BP3-depleted cells compared to WT MDA-MB-231 cells, which was reduced by rescue with WT HS1BP3 but not by the PRR3.1-mutant (Fig 6F-G). The endogenous TKS5 puncta were cytoplasmic and did not appear in F-actin-rich puncta as observed by phalloidin stain (Fig 6F), suggesting they were different from invadopodia. We also observed the presence of TKS5 signal in 10% of LAMP1-labeled late endosomes/lysosomes (Fig 6H). Importantly, HS1BP3 knockdown significantly increased the percentage of LAMP1-positive structures containing TKS5 in WT cells and in cells expressing the HS1BP3 PRR3.1-mutant (Fig 6H). The TKS5-positive LAMP1 structures were significantly closer to the nucleus in the PRR3.1-mutant cells (Fig 6I), which may suggest a buildup of TKS5-positive multivesicular endosomes that fail to secrete. Taken together, our data suggest that HS1BP3, and its interaction with cortactin, modulate the intracellular sorting and secretion of TKS5 and, by extension, cell invasiveness.

## Discussion

Here we demonstrate that an interaction between HS1BP3 and cortactin modulates cell proliferation and invasion, providing an explanation to the observed correlation between high HS1BP3 levels and poor survival of gastric adenocarcinoma and TNBC patients. Gastric cancer is currently the 5^th^ most common cancer type and is attributable to 8.3% of cancer deaths worldwide (34–36). Most gastric cancer patients are only diagnosed when the cancer has reached an advanced stage, and, with few effective treatments other than surgery, the prognosis remains poor (37). Similarly, breast cancer is the topmost cause of cancer death in women (38) with the TNBC subtype being the most lethal with no viable treatments (29). Hence, it is important to investigate the molecular mechanisms underlying these diseases for earlier detection and development of new responsive therapies.

We have previously shown that HS1BP3 functions as a negative regulator of autophagy and that it interacts with cortactin (6–8). To elucidate the molecular mechanisms underlying the negative correlation between HS1BP3 mRNA levels and survival time in gastric adenocarcinoma and in TNBC patients, we first pinpointed the interaction site between HS1BP3 and cortactin and generated an HS1BP3 mutant (PRR3.1) that abolished the interaction. Intriguingly, we demonstrated that the HS1BP3-cortactin interaction is important for efficient extracellular matrix degradation in the invasive TNBC cell line MDA-MB-231 and for cell proliferation in the non-invasive gastric adenocarcinoma cell line MKN-74.

High cortactin levels have previously been associated with worse cancer phenotypes (16–18,39,40), but there were no significant differences in the survival of high and low-expressing *CTTN* gastric adenocarcinoma and TNBC patients. Thus, it is possible that cortactin may not be transcriptionally regulated but has a function through its interaction to HS1BP3 (7). Comparing pairs of control and tumour tissue from gastric adenocarcinoma patients we found that *HS1BP3* is specifically upregulated in the tumour tissue, suggesting that *HS1BP3* is transcriptionally upregulated upon tumorigenesis. To our surprise, we did not detect any significant differences in basal- or starvation-induced autophagy in HS1BP3 KO cells compared to WT MKN-74 cells, suggesting that the effect of HS1BP3 on proliferation and invasion is independent of autophagy. Although, we have previously identified HS1BP3 as a negative autophagy regulator in U2OS, HeLa and HEK293 cells (6–8) and *in vivo* in Zebrafish (7), autophagy flux in metastatic cancer sites, from where MKN-74 cells are isolated, may be regulated differently.

To further understand how the HS1BP3-cortactin interaction modulates matrix degradation and proliferation, we investigated whether the expression level and/or cellular localization of the cortactin-binding protein TKS5 was affected in cells expressing the HS1BP3 PRR3.1 mutant. This was based on the observed co-expression between *HS1BP3* and *SH3PXD2A*, the gene encoding TKS5 in gastric tissues. TKS5 is a known interactor of cortactin in the initiation of invadopodia required to rearrange the cytoskeleton for invadopodia and podosome formation and invasion of the extracellular matrix (41,14). Loss of HS1BP3 had no effect on the interaction between TKS5 and cortactin, suggesting that HS1BP3 KO cells still can initiate invadopodia formation. However, we observed bright TKS5-GFP puncta significantly accumulating in cells expressing the HS1BP3 PRR3.1 mutant in both MKN-74 and MDA-MB-231 cells. We further demonstrated that these puncta, as well as endogenous TKS5 puncta, are LAMP1-positive. It was recently suggested that TKS5 is degraded by autophagy due to the observation of TKS5-GFP in LAMP1-structures (14), but as the low pH of the endolysosomes would have quenched the GFP-signal, it is more likely that TKS5 in LAMP1-structures is found in vesicles inside multivesicular bodies. Moreover, we did not observe an accumulation of TKS5 when treating MKN-74 cells with the lysosomal inhibitor Bafilomycin A1, suggesting that TKS5 is not a cargo for autophagy. Furthermore, there was no significant effect on autophagy levels when knocking down TKS5 (6,8). We propose that TKS5 in LAMP1-structures are inside multivesicular bodies destined for secretion.

In line with a buildup of TKS5 in multivesicular bodies in HS1BP3 PRR3.1 mutant cells, cortactin knockdown has been associated with reduced secretion and lost matrix degradation (18,17,40,16,39). Hence, inhibition of the HS1BP3-cortactin interaction may reduce matrix degradation by 1) a failure to initiate invadopodia due to increased endolysosomal localization of TKS5 and/or 2) reduced secretion of autocrine factors important for degradation. The effect of the HS1BP3-cortactin interaction on cell proliferation may also be explained by reduced secretion in the HS1BP3 PRR3.1 cells. Weaver et al (39) demonstrated that secreted factors from cortactin-overexpressed cells can rescue reduced proliferation in cortactin-depleted cells, suggesting that cortactin facilitates secretion of autocrine factors important for proliferation. It is not clear if TKS5 secretion is responsible for the proliferation and invasion effects observed in HS1BP3 PRR3.1 cells, but endosomal TKS5 trafficking (32) and secretion (25) has been found to be important for tumour growth and invasiveness. Integrating HS1BP3 and cortactin, we have previously shown that HS1BP3 localises on recycling endosomes (7), which is also the case for cortactin (15). Hence, it is possible that the HS1BP3-cortactin interaction regulates cancer through divergence of the endocytic pathway and regulation of secretion of TKS5 and/or or other autocrine factors important for cancer malignancy.

## Materials and Methods

### Reagents and Tools Table

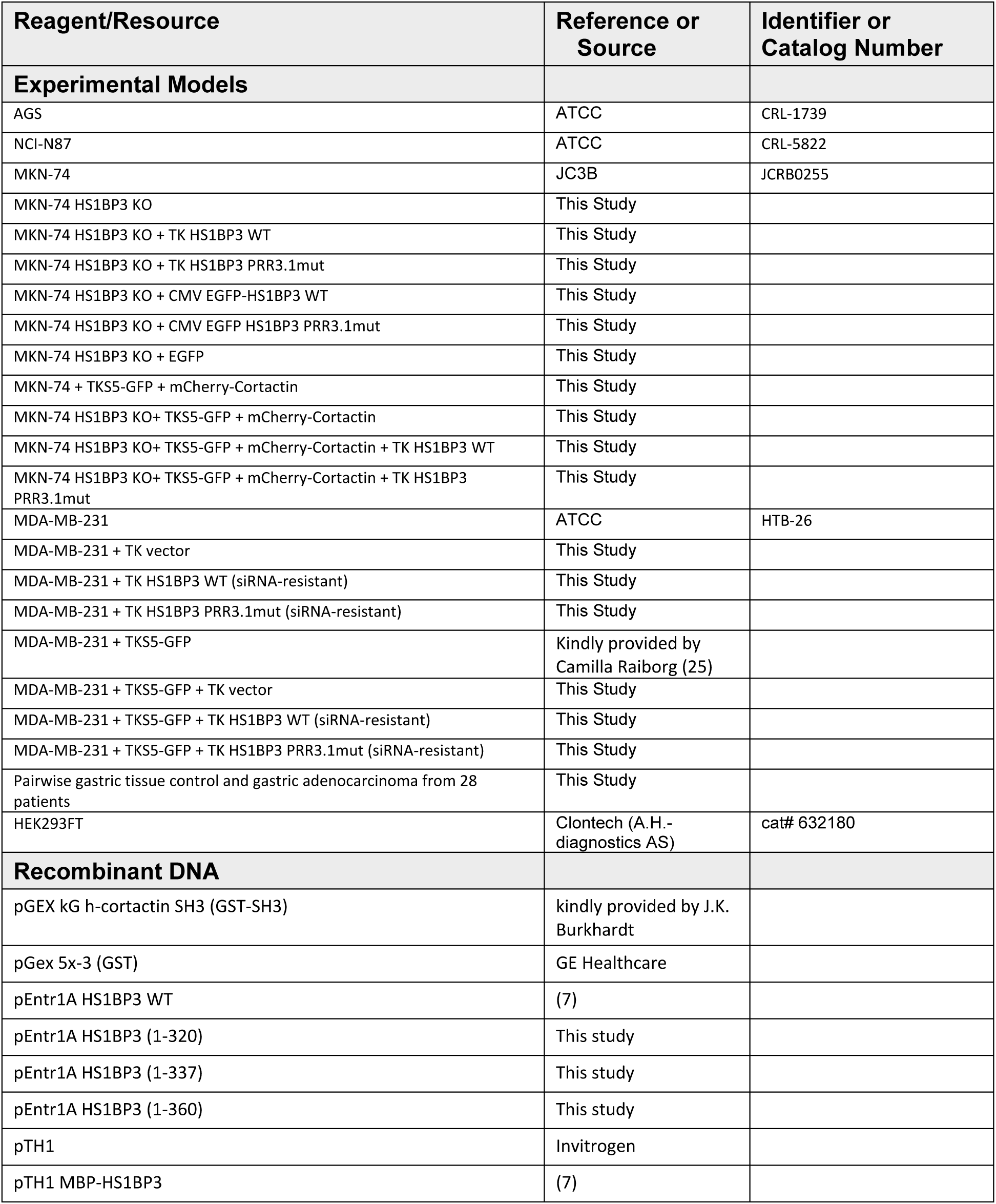
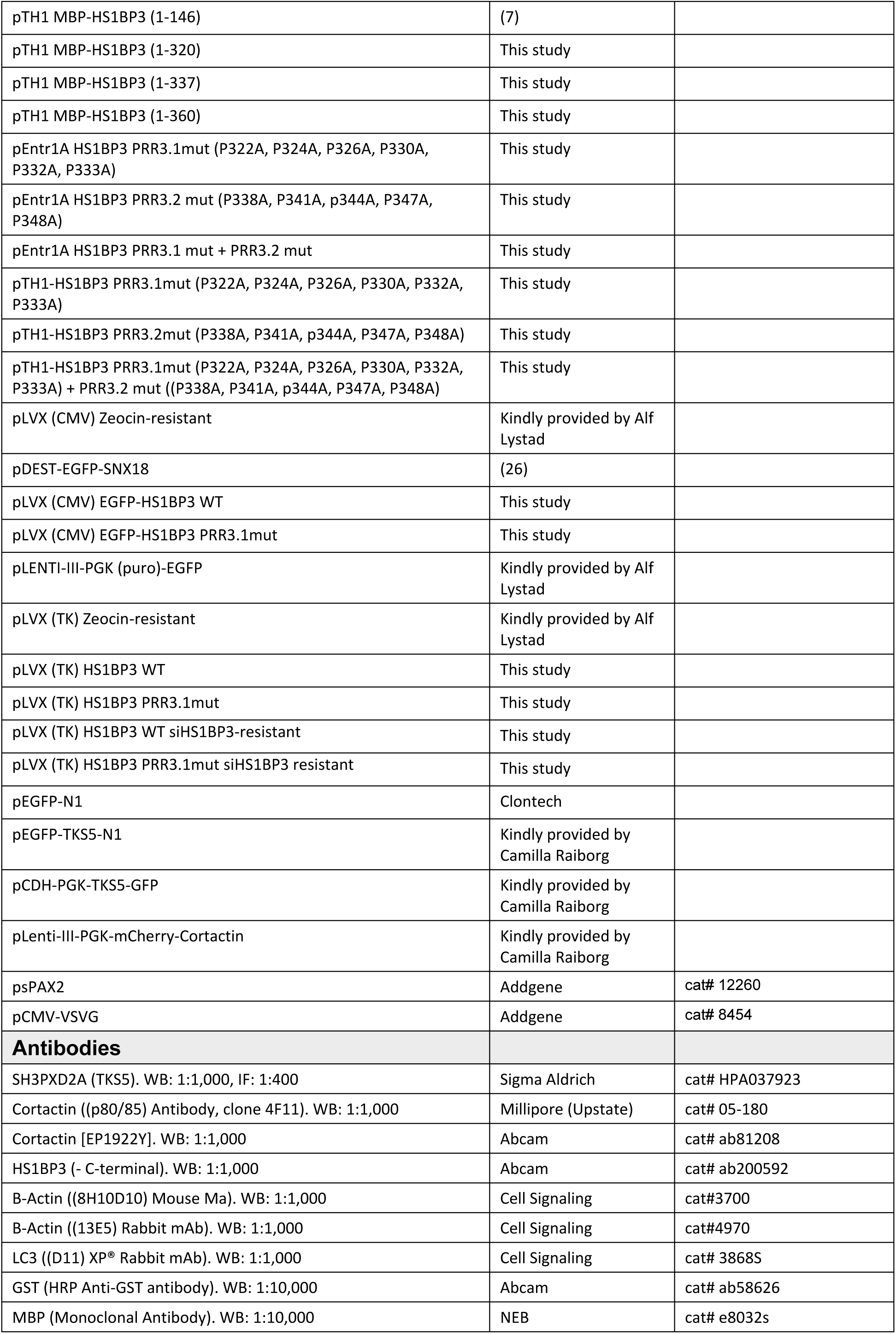
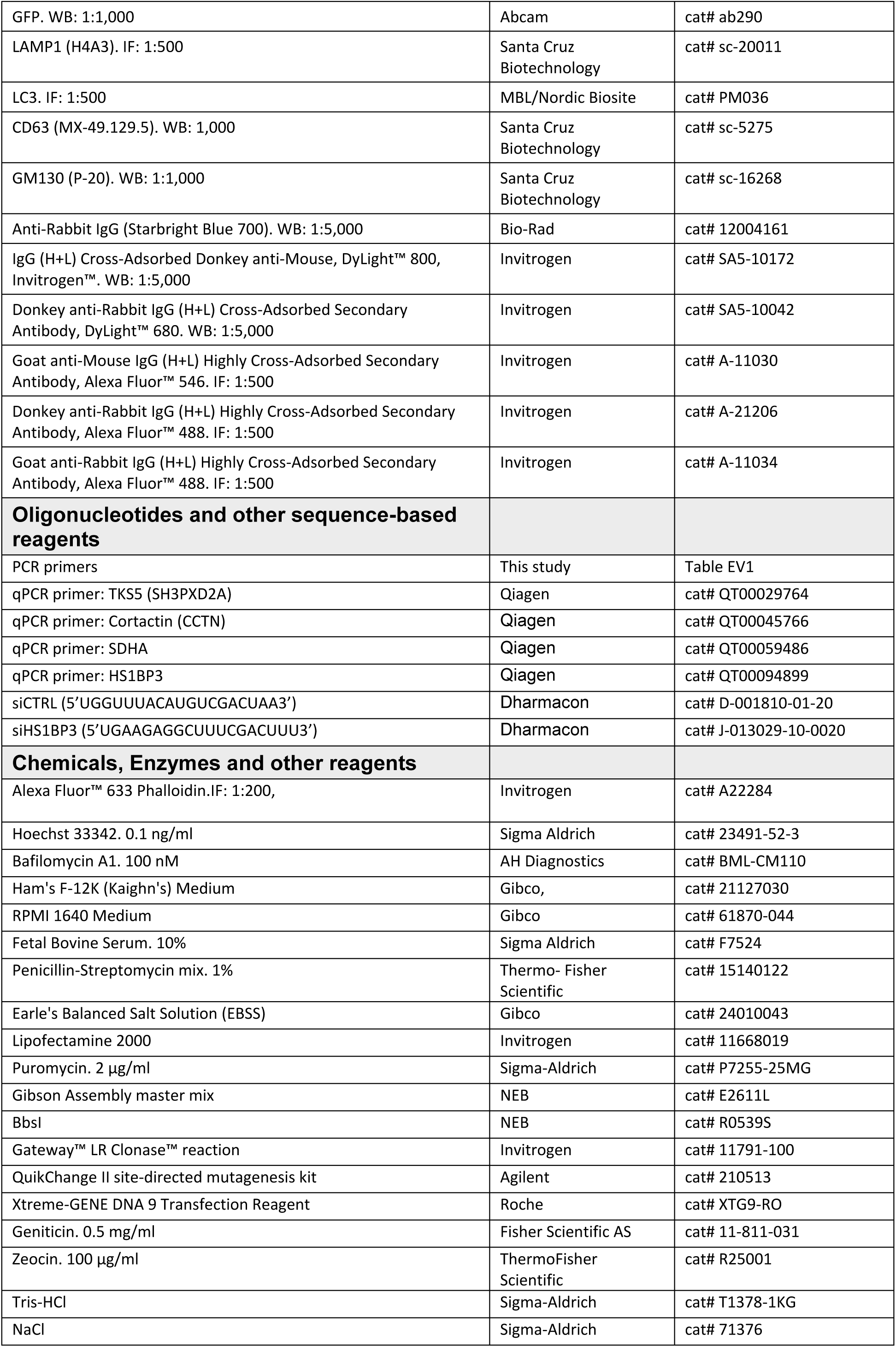
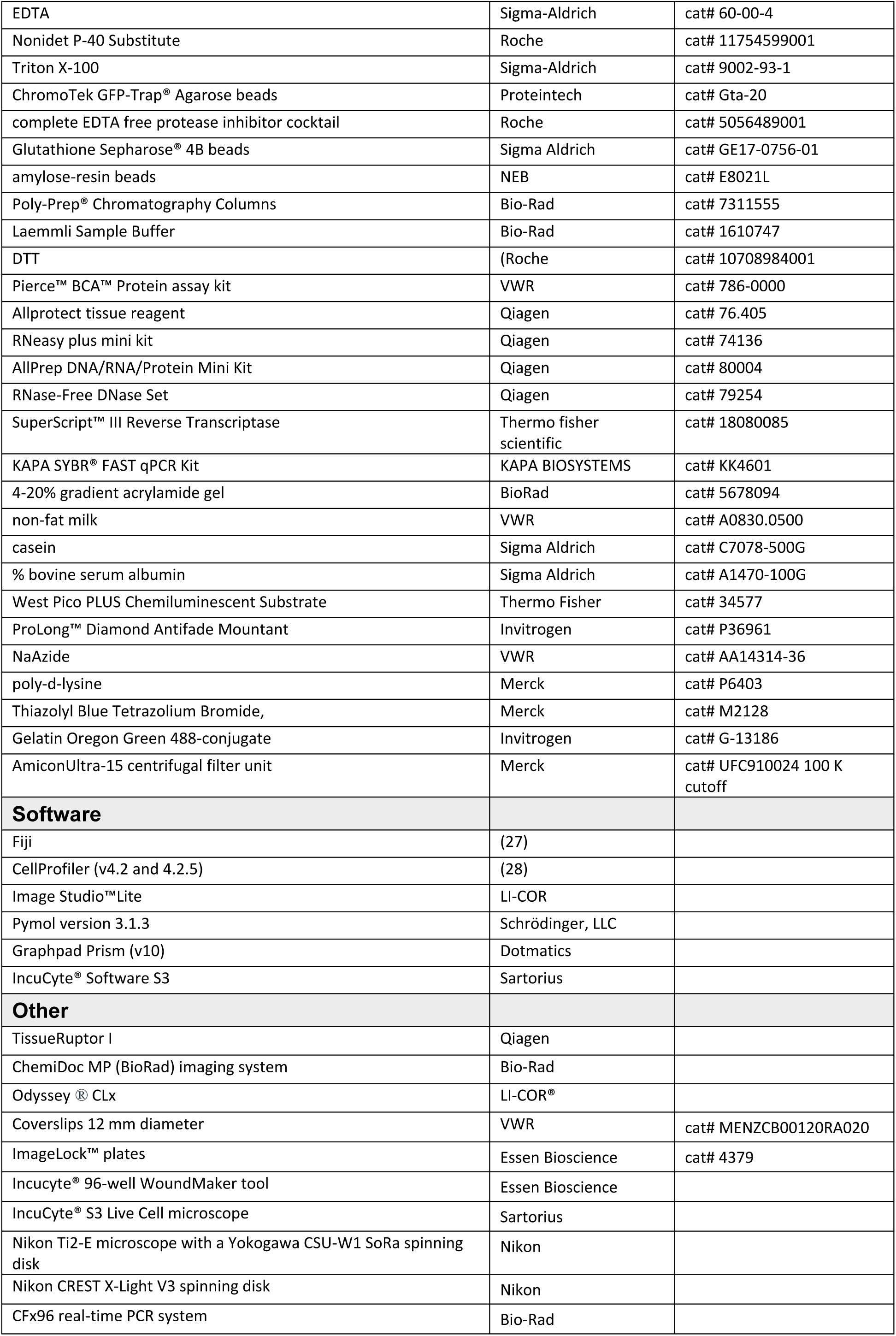

### Methods and Protocols

#### KM plot analysis

Kaplan-Meier plot analysis and Cox proportional hazards regression on overall survival or relapse-free survival are based on mRNA gene chip data from breast, ovarian, lung, gastric, colon, AML, and myeloma cancer patients and was performed using the KM-plotter database and interface www.kmplot.com (21,22). Comparison groups were split according to high (third quartile) and low (first quartile) levels of the gene of interest by trichotomization. For the effect of HS1BP3 expression, indicated Hazard ratios with 95% confidence intervals and p-values were plotted on a forest plot for comparison on effect of HS1BP3 in different cancers.

#### Gene expression set analysis

To study gene expression data of healthy and cancerous tissues, Xena (23) was used to filter GTex Portal and TCGA data for the gene(s) and tissue(s) of interest.

#### AlphaFold model

The interaction between the full-length HS1BP3 (Q53T59) and the SH3 fragment of cortactin (Q14247: amin acids 492-550) was modelled in AlphaFold Server (AlphaFold V3) https://alphafoldserver.com/ (24). All five ranked models were investigated using Pymol version 3.1.3 (Schrödinger, LLC) where all models had similar interactions and one is shown.

The InterfaceResidues script https://www.pymolwiki.org/index.php/InterfaceResidues was used to illustrate interacting residues in cloud view and the Konstantin Korotkov code was used to show AlphaFold pLDDT colour scheme https://github.com/sokrypton/ColabFold.

#### Cell culture

AGS (CRL-1739), NCI-N87 (CRL-5822) and MDA-MB-231 (HTB-26) were purchased from ATCC and MKN-74 (JCRB0255) was purchased from JC3B cell bank. AGS were cultured in Ham’s F-12K (Kaighn’s) Medium (Gibco, 21127030) supplemented with 10% Fetal Bovine Serum (FBS; Sigma Aldrich, #F7524) and 1% penicillin-Streptomycin mix (Thermo-Fisher Scientific, #15140122) while MDA-MB-231, MKN-74 and NCI-N87 cells were maintained in RPMI 1640 Medium (Gibco, 61870-044) in 10% FBS and 1% PS. All cells were cultured in a humidified incubator at 37 °C with 5% CO_2_. MKN-74 and MDA-MB-231 cells with stable overexpression of HS1BP3 were kept in the indicated medium with appropriate resistance antibiotics. To starve the cells in nutrient-deplete medium, they were incubated in Earle’s Balanced Salt Solution (EBSS; Gibco, 24010043). MDA-MB-231 WT cells and MDA-MB-231 cells stably overexpressing TKS5-GFP were a kind gift from C. Raiborg (25). All cells used in this study have been testing negative for mycoplasma in the past three years.

#### Generation of MKN-74 HS1BP3 KO

To generate MKN-74 HS1BP3 KO cells, CRISPR/Cas9 KO plasmids (px459) were digested with BbSI and forward and reverse guide RNAs 1 (fw 5’CACCGCGCCGTCATGCAGTCCCCGG3’, rv 5’AAAC CCGGGGACTGCATGACGGCG3’) and 2 (fw 5’CACCgCCCCAGCACCAGGAGGTACG3’, rv 5’AAACCGTACCTCCTGGTGCTGGGGc3’) were annealed respectively, BbSI-digested and ligated into the digested px459 plasmid. MKN-74 cells were transfected with the CRISPR/Cas9 KO plasmids using Lipofectamine 2000 (Invitrogen, 11668019) and subsequently selected for positive clones for maximum 48 hours with 2 µg/ml puromycin (Sigma-Aldrich, P7255-25MG).

#### Plasmids

All plasmids were made using Gibson Assembly master mix (NEB, E2611L), Gateway™ LR Clonase™ reaction (Invitrogen, 11791-100) or the QuikChange II site-directed mutagenesis kit (Agilent, 210513) using primers as indicated in Extended View Table I.

#### Generation of stably overexpressing cells

MKN-74 WT and HS1BP3 KO cells and MDA-MB-231 cells with stable overexpression were made by co-transfecting indicated expression plasmids with psPAX2 and pCMV-VSVG at 1:1:1 ratio using Xtreme-GENE DNA 9 Transfection Reagent (Roche, XTG9-RO) into HEK293FT cells to produce lentiviral particles that were transduced into target cells. Positive cells were selected with 1 µg/ml puromycin (Sigma-Aldrich, P7255-25MG), 0.5 mg/ml Geniticin (Fisher Scientific AS, 11-811-031) or 100 µg/ml zeocin (ThermoFisher Scientific, R25001) depending on the antibiotic resistance of the transduced expression plasmid.

#### Transient overexpression of EGFP and TKS5-EGFP

For transient transfection of EGFP (pEGFP-N1) and TKS5-EGFP (pEGFP-TKS5-N1), 8.0x10^5^ MKN-74 cells were seeded into 10 cm dishes and left for 48 hours before transfection of the indicated plasmids at equimolar concentrations of each gene using lipofectamine 2000 (Invitrogen, 11668019).

#### siRNA knockdown

Cells were incubated with Dharmacon oligonucleotides targeting each gene at 20 nM, in combination with RNAiMAX (Invitrogen, 13778150) through reverse transfection for 72 hours. The siRNA nucleotides used were: siCTRL (5’UGGUUUACAUGUCGACUAA3’, #D-001810-01-20), siHS1BP3 (5’UGAAGAGGCUUUCGACUUU3’, #J-013029-10-0020).

#### In vitro protein production and purification

GST- and MBP-tagged proteins were expressed in SoluBL21 *E.coli* cells. GST-bound proteins were purified by lysing the *E.coli* cells with GST-lysis buffer (50 mM Tris-HCl (pH8; Sigma-Aldrich, T1378-1KG), 250 mM NaCl (Sigma-Aldrich, 71376), 1 mM (Roche, 10708984001), 1x complete EDTA free protease inhibitor cocktail (Roche, 5056489001)) and isolating the proteins with Glutathione Sepharose® 4B beads (Sigma Aldrich, GE17-0756-01). MBP-tagged proteins were purified by lysing the *E. coli* cells in MBP-lysis buffer (20 mM Tris-HCl pH 7.4, 200 mM NaCl, 1 mM EDTA, 1 mM DTT, 1x complete EDTA free protease inhibitor cocktail) and isolating the proteins with amylose-resin beads (NEB, E8021L), followed by elution with 10 mM maltose on Poly-Prep® Chromatography Columns (Biorad, 7311555) at 4°C. For *in vitro* interaction assays, equal concentrations of GST- and MBP-tagged proteins were incubated together for 2 hours at 4°C and washed 6 times with GST-buffer before heat inactivation and denaturation at 95°C in Laemmli Sample Buffer (Bio-Rad, 1610747) supplemented with DTT (Roche, 10708984001) followed by immunoblotting. The same procedure was done for cell lysate incubation with GST-tagged proteins, here 450 µg protein lysate (as measured using Pierce™ BCA™ Protein assay kit (VWR, 786-0000)) was added.

#### Cell lysis and immunoprecipitation

Cells were lysed in RIPA buffer (50 mM Tris-HCl (pH 7.5; Sigma-Aldrich, T1378-1KG), 150 mM NaCl (Sigma-Aldrich, #71376), 1 mM EDTA (Sigma-Aldrich, #60-00-4), 0.05% Nonidet P-40 Substitute (Roche, 11754599001), 0.25% Triton X-100 (Sigma-Aldrich, #9002-93-1)) and centrifuged at 15,000xg for 10 min. The protein concentration of the soluble protein fraction was measured using the Pierce™ BCA™ Protein assay kit (VWR, 786-0000). For GFP-trap assays, at least 450 µg protein lysate or the whole amount of lysate was added to 15 µl homogenized and pre-cleared ChromoTek GFP-Trap® Agarose beads (Proteintech, Gta-20) for 2.5 hours before washing the beads four times in RIPA buffer before heat inactivation and denaturation at 95°C in Laemmli Sample Buffer (Bio-Rad, 1610747) supplemented with DTT (Roche, 10708984001) before immunoblotting.

#### Patient sample storage and handling

Biopsies were obtained from 28 patients diagnosed with gastric adenocarcinoma at Akershus University Hospital, Norway. On admission for surgery, each participant gave a written, informed consent to participate in the study. Procedures conducted included complete gastrectomy or partial gastrectomy depending on the tumour location in the stomach. The biopsies were taken within 10 minutes after removal of the surgical specimen. From each patient, two tumour samples as well two control samples from healthy gastric tissue adjacent to the tumour were obtained. One set of tumour and control samples were directly placed in Allprotect tissue reagent (Qiagen, 76.405) for long-term storage at –20°C, and the other set was directly stored in 10% formalin solution for immunohistochemistry analysis and placed at 4 °C. After 24 hours, the formalin was diluted to 1% for prolonged storage at 4 °C. RNA extraction of Allprotect samples is described below.

#### RNA isolation and qPCR

Cells were trypsinised and washed in PBS before RNA extraction using the RNeasy plus mini kit (Qiagen,74136) according to the manufacturer’s instructions. To extract RNA from the Allprotect-stored bioposies, the AllPrep DNA/RNA/Protein Mini Kit (Qiagen, 80004) was used according to the manufacturer protocol with TissueRuptor I (Qiagen) homogenisation of the tissue and additional on-column DNA digestion with the RNase-Free DNase Set (Qiagen, 79254). The RNA was reverse transcribed using the SuperScript™ III Reverse Transcriptase (Thermo fisher scientific, 18080085) and qPCR was performed using KAPA SYBR® FAST qPCR Kit (KAPA BIOSYSTEMS, KK4601) in a CFx96 real-time PCR system (Bio-Rad) with Qiagen QuantiTect primers (Qiagen, 249900). Transcript levels of genes of interest were normalised to SDHA and fold change quantifications were performed using the 2^−ΔΔCt^ method.

#### Protein extraction and immunoblotting

Cells were lysed in RIPA buffer (either: 50 mM Tris-HCl (pH 7.5; Sigma-Aldrich, T1378-1KG), 150 mM NaCl (Sigma-Aldrich, #71376), 1 mM EDTA (Sigma-Aldrich, #60-00-4), 0.05% Nonidet P-40 Substitute (Roche, 11754599001), 0.25% Triton X-100 (Sigma-Aldrich, #9002-93-1); or: 20 mM Tris-HCL (pH 7.4), 150 mM NaCl, 5 mM EDTA and 1% Triton X-100) and protein concentration was measured using the Pierce™ BCA™ Protein assay kit (VWR, 786-0000) and denatured at 95°C in Laemmli Sample Buffer (Bio-Rad, 1610747) supplemented with DTT (Roche, 10708984001). At least 20 µg of protein was loaded on 4-20% gradient acrylamide gel (BioRad, 5678094) and transferred to a PVDF membrane. For Ponceau S staining, the membrane was dried and re-wetted with 100% methanol before Ponceau S staining followed by membrane drying and imaging in the ChemiDoc MP (BioRad) imaging system. The membranes were blocked in either 5% non-fat milk (VWR, A0830.0500) or 1% casein (Sigma Aldrich, C7078-500G) in 1X PBS for a minimum of 45 minutes before primary antibody incubation in 5% bovine serum albumin (Sigma Aldrich, A1470-100G) in 1X PBS overnight at 4°C. The membranes were washed and incubated with secondary antibody diluted in 1X PBS for a minimum of 45 min at room temperature before washing and imaging. For infrared fluorescent signal, the Odyssey ® CLx (LI-COR®) and ChemiDoc MP were used to image the membranes. For chemiluminescent signal, the SuperSignal West Pico PLUS Chemiluminescent Substrate (Thermo Fisher, 34577) was used according to manufacturer instructions and the signal detected in the ChemiDoc MP. Band quantifications were done in Image Studio™Lite and Fiji (27).

#### Immunofluorescence

For immunofluorescence, cells in complete growth medium were fixed by directly adding 2x 4% PFA-PHEM buffer and incubating the cells for 15 min at 37°C with 1:10,000 Hoechst 33342 (Sigma Aldrich, 1 µg/ml). Cells were then washed 3x in PBS and incubated with primary antibody diluted in 0.05% saponin in PBS-BSA 3% for 1-2 hours. After three washes in PBS, the cells were incubated with the corresponding secondary antibodies diluted in PBS for 40 minutes-1 hour and finally washed 3x in PBS. If cells were grown on coverslips, the coverslips were mounted in ProLong™ Diamond Antifade Mountant (Invitrogen, P36961). If grown in 8-well chambers, the cells were kept in PBS with 0.02% NaAzide (VWR, AA14314-36) at 4°C. Image analysis was carried out using CellProfiler software (v4.2 and 4.2.5) (Stirling et al., 2021). The images were separated into individual channels and in some experiments enhanced to improve the visibility of key features. Nuclei, cells, and puncta were detected by applying a manual threshold to segment the areas of interest. Co-localisation was assessed by relating the structures.

For MKN-74 cells, the glass coverslip/8-well chamber bottom was coated with poly-d-lysine (Merck, P6403) for minimum 30 min at 37°C and washed 1x with PBS before cell seeding to ensure attachment.

#### Wound healing assay

Cells were seeded to confluency in 96 well ImageLock™ plates (Essen Bioscience, 4379) and left overnight. To make wounds, the Incucyte® 96-well WoundMaker tool (Essen Bioscience) was used and the medium changed before the plate was left for up to 80 hours in the IncuCyte® S3 Live Cell microscope (Sartorius) for imaging every 10 min. Quantification of relative wound density (compared to time 0 hour) was done using the IncuCyte® Software S3 (Sartorius) by measuring the “spatial cell density” in the wound area relative to the spatial cell density outside of the wound area at every time point.

#### Cell Viability assay

Cells were seeded to 70% confluence in 96-well plates and left overnight before adding MTT viability reagent (Thiazolyl Blue Tetrazolium Bromide, Merck, M2128) to a working concentration of 0.5 mg/mL for four hours at 37°C in humidified chamber. Finally, the liquid was removed, and the resulting formazan crystals were dissolved in DMSO before measuring absorbance at 570 nm.

#### Proliferation assays

Cells were seeded to 2-10% confluency in 96-well plates and left for 51 hours in IncuCyte® S3 Live Cell microscope (Sartorius) with imaging every 1.5 hour. Confluence was quantified in the IncuCyte®

Software S3 (Sartorius) and data presented as relative confluence compared to time 0 using the mean of three replicate wells from three independent experiments.

#### Gelatin degradation assay

To prepare gelatin-coated coverslips, 12 mm diameter (VWR, MENZCB00120RA020) coverslips were sterilised by laying flat on parafilm in laminar flow cell hood with UV light treatment for 60 min. Then, while continuing to work in the cell hood, 30 µl preheated (37°C) 0.2 mg/ml Gelatin Oregon Green 488-conjugate (Invitrogen, G-13186) in 2% sucrose in PBS was dropped on clean parafilm and the sterilised coverslips left on top for 20 min at room temperature in the dark. 20 ul of 0.5% glutaraldehyde in PBS was left on a clean sheet of parafilm and the gelatin-coated coverslips moved on top gelatin-side down. The coated coverslips were fixed for minimum 40 min followed by 3x wash in PBS and kept in 1% penicillin-streptividin in PBS at 4 degrees until use. Before use, the coverslips were incubated in complete cell culture medium for at least 60 min before cell seeding cells on top for the indicated time and subsequently fixed in 4% paraformaldehyde in PHEM for 15 min at 37°C. The cells were subsequently permeabilised in 0.05% saponin and stained in 1:200 solution of Phalloidin-633 (Invitrogen, A22284) and 1:10,000 Hoechst 33342 (Sigma Aldrich, 1 µg/ml) in 0.05% saponin in PBS for 40 min before 3x wash in PBS and mounting in ProLong™ Diamond Antifade Mountant (Invitrogen, P36961). The number of cells covering degradative areas in the geleatin (completely black spots) were counted and compared relative to total number of cells observed.

#### Extracellular vesicle enrichment

Extracellular vesicle enrichment using 100 K spin-column filtration was done as described previously (25). Briefly, 4 × 10^6^ cells were seeded in 150 mm dishes and incubated for 48 hours before washing the dishes 2x in 12 ml serum-free RPMI (Gibco, 61870-044) followed by incubation for 44 hours in 14 ml serum-free RPMI. Then the conditioned media was centrifuged at 300 × g at 4 °C for 10 min, at 1000 × g at 4 °C for 10 min and at 10,000 × g at 4 °C for 30 min. This was followed by concentration of extracellular vesicle fraction to about 300 µl using an AmiconUltra-15 centrifugal filter unit (UFC910024 100 K cutoff) at 4,000 × g for 8 minutes. The cells were trypsinised and counted and 10% of the cell lysate and 8% of the concentrated media was used for analysis by western blot.

### Statistical analysis

Statistical analysis and preparation of figures, except for the KM-analysis, was performed using Graphpad Prism with tests as indicated in the figure legends.

### Ethical considerations

This project has been under the necessary ethical considerations and the collection, storage and use of gastric tissue from gastric adenocarcinoma patients was pre-approved by the Norwegian Regional committee for medical and health research ethics (reference #166245).

## Acknowledgements

We would like to acknowledge the past and current members of the Simonsen lab for critical discussions and feedback, particularly Anette Dahl who helped with AlphaFold and Pymol analysis, Alf Lystad who shared plasmids and Santosh Phuyal for sharing the method for gelatin-coating of coverslips. A special thanks goes to Nina-Marie Pedersen, Camilla Raiborg and Eva Wenzel in the Stenmark group for consultation, critical review and sharing of resources and methods.

The IncuCyte S3 imaging was performed at two core facilites: Cancer Immunology Department at the Institute for Cancer Research, The Norwegian Radium Hospital and; at the MolMed Imaging Platform at the Institute of Basic Medical Sciences, University of Oslo. The confocal imaging was performed at the Advanced Light Microscopy Core Facility at the Institute for Cancer Research, The Norwegian Radium Hospital. We acknowledge the Norwegian Core Facility for Human Pluripotent Stem Cells at the Norwegian Center for Stem Cell Research for mycoplasma testing.

This project was carried out with funding from the South-Eastern Norway Regional Health Authority (grant no. 2020032), the Norwegian Cancer Society (grant no. 190251) and the Research Council of Norway through its Centers of Excellence funding scheme (grant no. 262652).

## Author contributions

AAL: Conceptualization, formal analysis, investigation, methodology, project administration, validation, visualization, and writing—original draft, review, and editing

KS: Conceptualization, formal analysis, investigation, methodology, project administration, writing – review and editing

CV: Investigation, formal analysis, writing – review and editing LTM: Investigation, writing – review and editing

NA: Investigation, writing – review and editing RG: Resources, writing – review and editing HK: Investigation, writing – review and editing

LE: Resources, writing – review and editing

AS: Supervision, conceptualization, methodology, project administration, funding acquisition, writing – review and editing

## Data availability

This study includes no data deposited in external repositories.

## Abbreviations

Arp2/3: Actin-related protein 2/3 complex
CTTN: cortactin
F-actin: Filamentous actin
HR: Hazard Ratio
HS1BP3: HCLS1-binding protein 3
SH3: Src Homology domain 3
KM: Kaplan Meier
LAMP1: Lysosome associated membrane glycoprotein 1
LC3: microtubule-associated protein light-chain 3 protein
PRR: Proline Rich Region
TNBC: Triple Negative Breast Cancer.

---

**Figure EV1:**
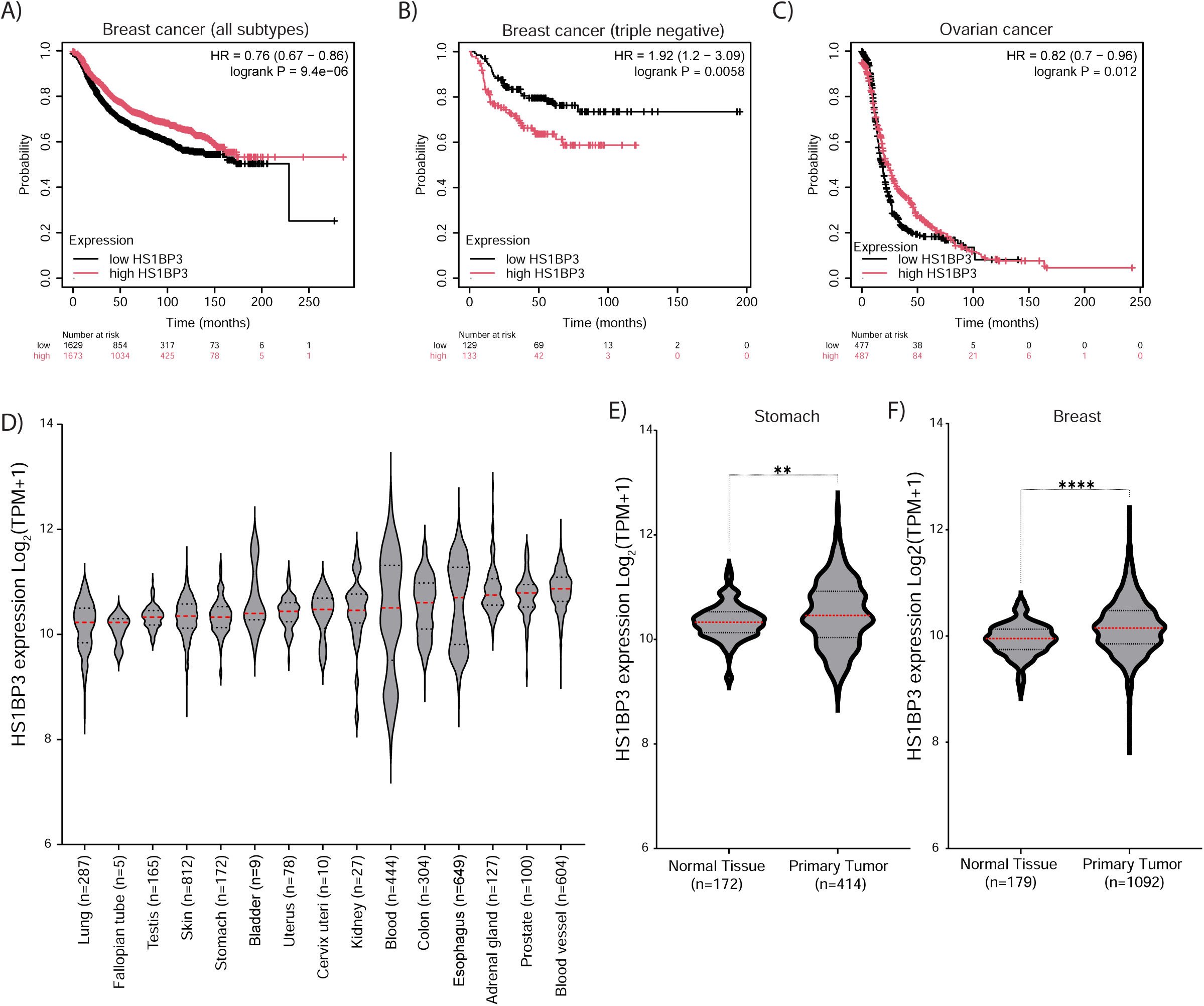
HS1BP3 levels is predictive for patient outcomes in breast and ovarian cancer. (**A-C**) Kaplan Meier plots showing the probability of survival in patients with breast cancer (all subtypes) (**A**), triple negative breast (**B**) and ovarian cancer (**C**) according to high (T3=third highest expressing tertile) and low (T1= third lowest expressing tertile) mRNA expression of HS1BP3 in tumour at time of detection. HR=Hazard ratio and p-value were determined with Cox proportional hazards regression. Plots are made in the KM plotter database. (**D**) Graph displaying the median and interquartile range of bulk tissue gene expression (TPM= transcripts per million + 1) of HS1BP3 across the top 15 most-HS1BP3 expressing human tissues according to median levels. The points are outliers more than 1.5x the interquartile range. The graph is extracted from GTex portal. (**E-F**) Graph displaying the median and interquartile range of bulk tissue gene expression of HS1BP3 in healthy (GTex portal database) and cancerous (TCGA database) tissues of gastric (**E**) and breast (**F**) p-values were determined using unpaired t-test. **= P <0.01; ****= P<0.0001.

**Figure EV2:**
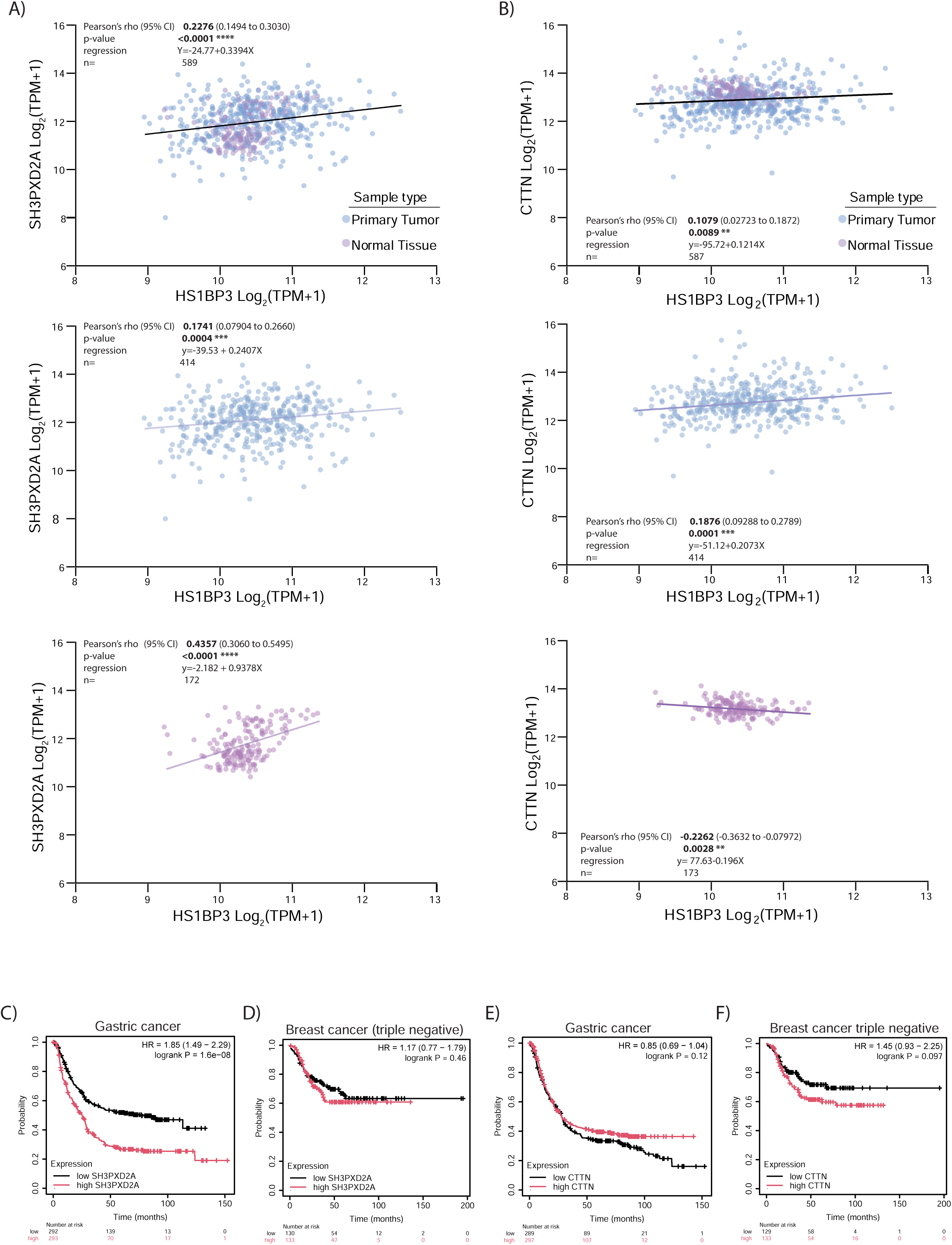
HS1BP3 and SH3PXD2A (TKS5) gene expressions correlate in gastric tissues. (**A-B**) Regression plots showing the Log_2_ gene transcripts per million+1 (TPM+1) of HS1BP3 correlated to the Log_2_ gene transcripts per million+1 (TPM+1) of SH3PXD2A (**A**) and CTTN (cortactin; **B**) in gastric tissues. The data is derived from TCGA (Gastric adenocarcinoma) and GTEx portal (healthy gastric tissue). A two-tailed Pearson’s correlation rho coefficient with 95% CI (confidence interval) and p-values are listed together with a simple linear regression line of best fit. (**C-F)** Kaplan Meier plots showing the probability of survival in patients with gastric (**C; E**) and triple negative breast cancer (**D; F**) according to high (T3=third highest expressing tertile) and low (T1= third lowest expressing tertile) mRNA expression of *SH3PXD2A* (**C**-**D**) or *CTTN* (**E-F**) in tumour at time of detection. HR=Hazard ratio and p-value were determined with Cox proportional hazards regression. Plots are made in the KM plotter database. **= P <0.01; ***= P <0.001; ****= P<0.0001.

**Figure EV3:**
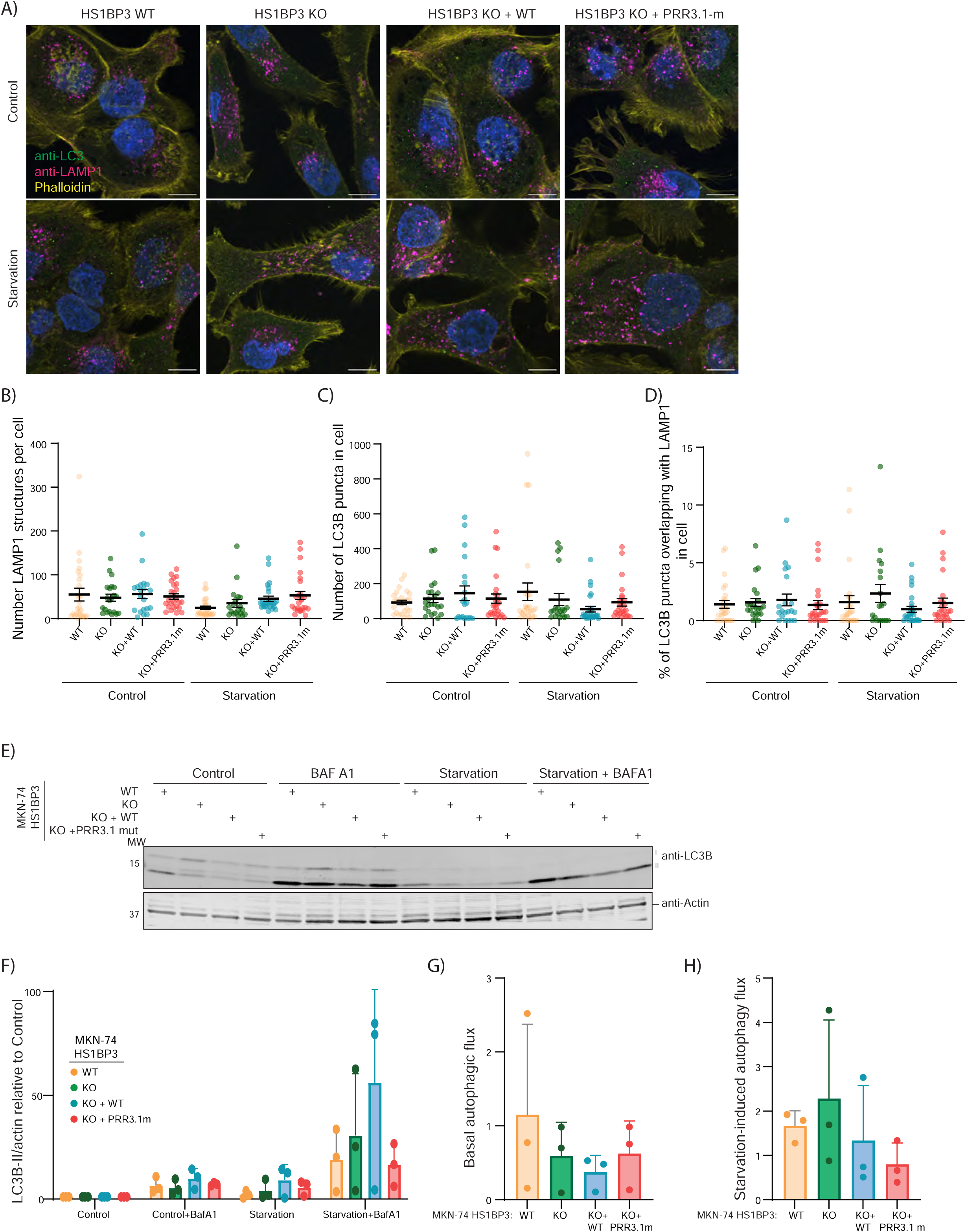
HS1BP3 KO or expression of the PRR3.1-mutant has no effect on autophagy in gastric adenocarcinoma. (**A**) Representative images of MKN-74 cells with HS1BP3 WT and KO 2.4 with HS1BP3 WT and PRR3.1 mutant. The cells were treated with EBSS for two hours for starvation, followed by immunofluorescence with antibodies anti-LC3B and anti-LAMP1 and staining with Phalloidin-633 and Hoechst, followed by acquisition with a Nikon Ti2-E microscope with a Yokogawa CSU-W1 SoRa spinning disk 100×/1.45 NA oil immersion objective. Scale bar: 10 µm. (**B**) Quantification of the number of LAMP1 structures per cell. (**C**) Quantification of the number of LC3B puncta per cell. (**D**) Quantification of the percentage of the overlap of LC3B with LAMP1 structures. Data are mean ± standard error of the mean with individual data points corresponding to a single field of view (n=3, 29-357 cells were quantified in each condition). (**E**) Representative immunoblots of the indicated antibodies in MKN-74 WT, HS1BP3 KO 2.4, and HS1BP3 KO 2.4 rescue with HS1BP3 WT and PRR3.1-mutant cell line treated with 2 hours starvation with and without 100 nM Bafilomycin A1 (BAFA1) to demonstrate the autophagy flux of the indicated cells, n=3. (**F**) quantification of (E) showing level of LC3B-II normalised to actin levels and relative to control condition of the indicated cell line. (**G**) Quantified basal autophagy flux from (E) measured as the level of LC3B-II in Control+BafA1 condition subtracted with the level of LC3B-II in control condition per cell line. (**H**) Quantified starvation-induced autophagy flux from (E) measured as level of LC3B-II in Starvation+BafA1 condition subtracted with level of LC3B-II in Starvation condition per cell line. (F-H) Data are mean ± Standard deviation of the mean with individual data points corresponding to each replicate. The statistical significance was calculated with ordinary one-way ANOVA followed by Tukey’s multiple comparison test (B-D), two-way ANOVA with Dunnett’s multiple comparison test (F), or one-way ANOVA with Dunnett’s multiple comparison test (G-H). Non-significant differences are not depicted.

**Figure EV4:**
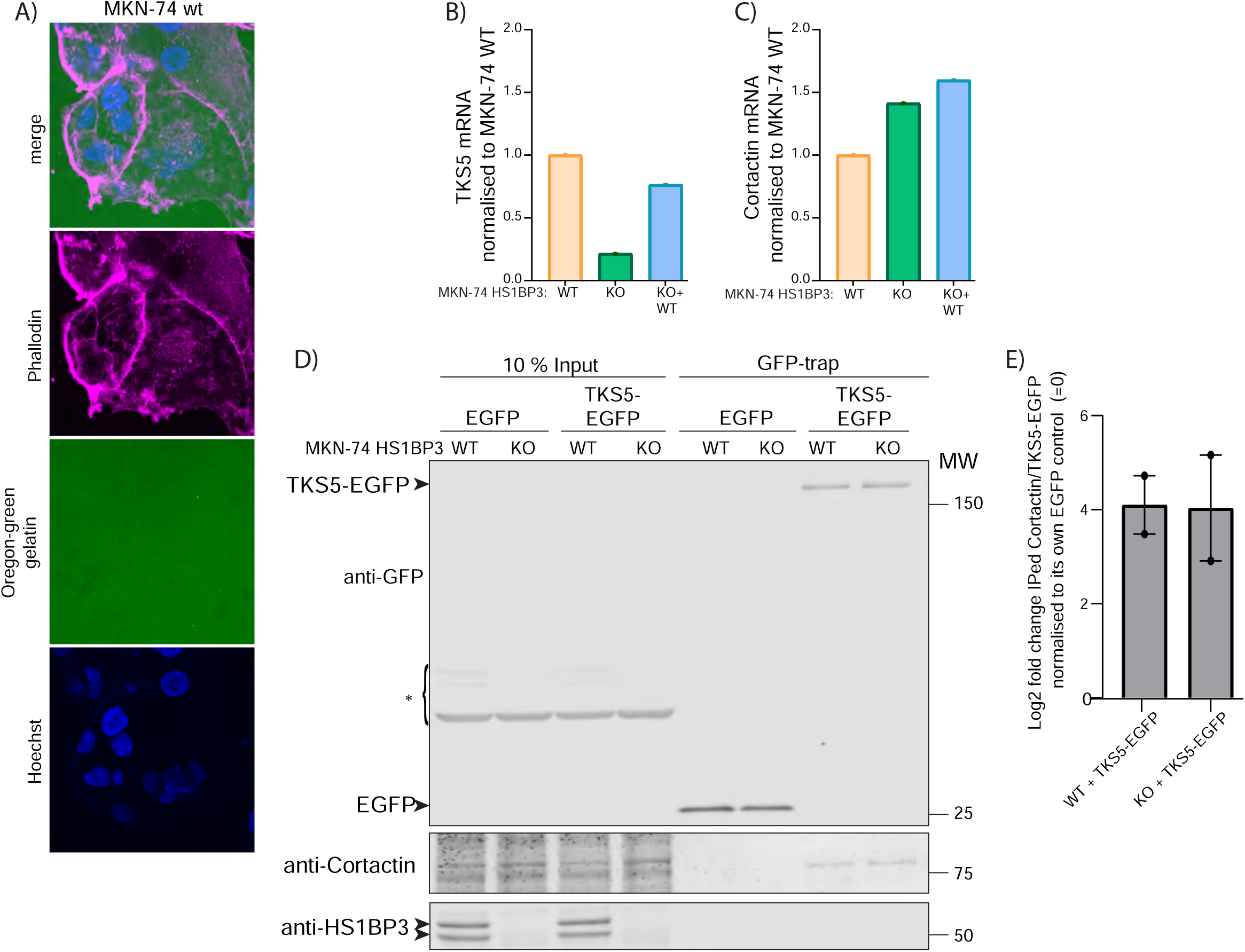
HS1BP3 levels do not affect cortactin-TKS5 protein interaction. (**A**) representative immunofluorescence image of MKN-74 seeded on coverslips coated with Oregon-Green labelled gelatin for 17 hours before PFA-fixation and staining with Hoechst and phalloidin before visualising the cells by confocal imaging with a Yokogawa CSU-W1 SoRa spinning disk 60xWI water immersion objective. (**B-C**) The mRNA expression of *SH3PXD2A* (TKS5) (B) and *CTTN* (cortactin) (C) was analysed by quantitative PCR. N=1. (**D**) MKN-74 WT and HS1BP3 KO (2.4) cells transiently transfected with pEGFP-N1 and pEGFP-TKS5-N1 were subjected to GFP-trap and immunoblot analysis of the indicated proteins. *=unspecific bands (**E**) quantification of (D) showing Log2 fold difference in mean level of cortactin compared to level of immunoprecipitated TKS5-EGFP. The data is normalised to level of cortactin compared to level of immunoprecipitated EGFP in each cell type (=0, not shown). The error bars are standard deviation, n=2. Differences between the conditions were not significant and determined by paired t-test (E).

**Extended View Table I.**
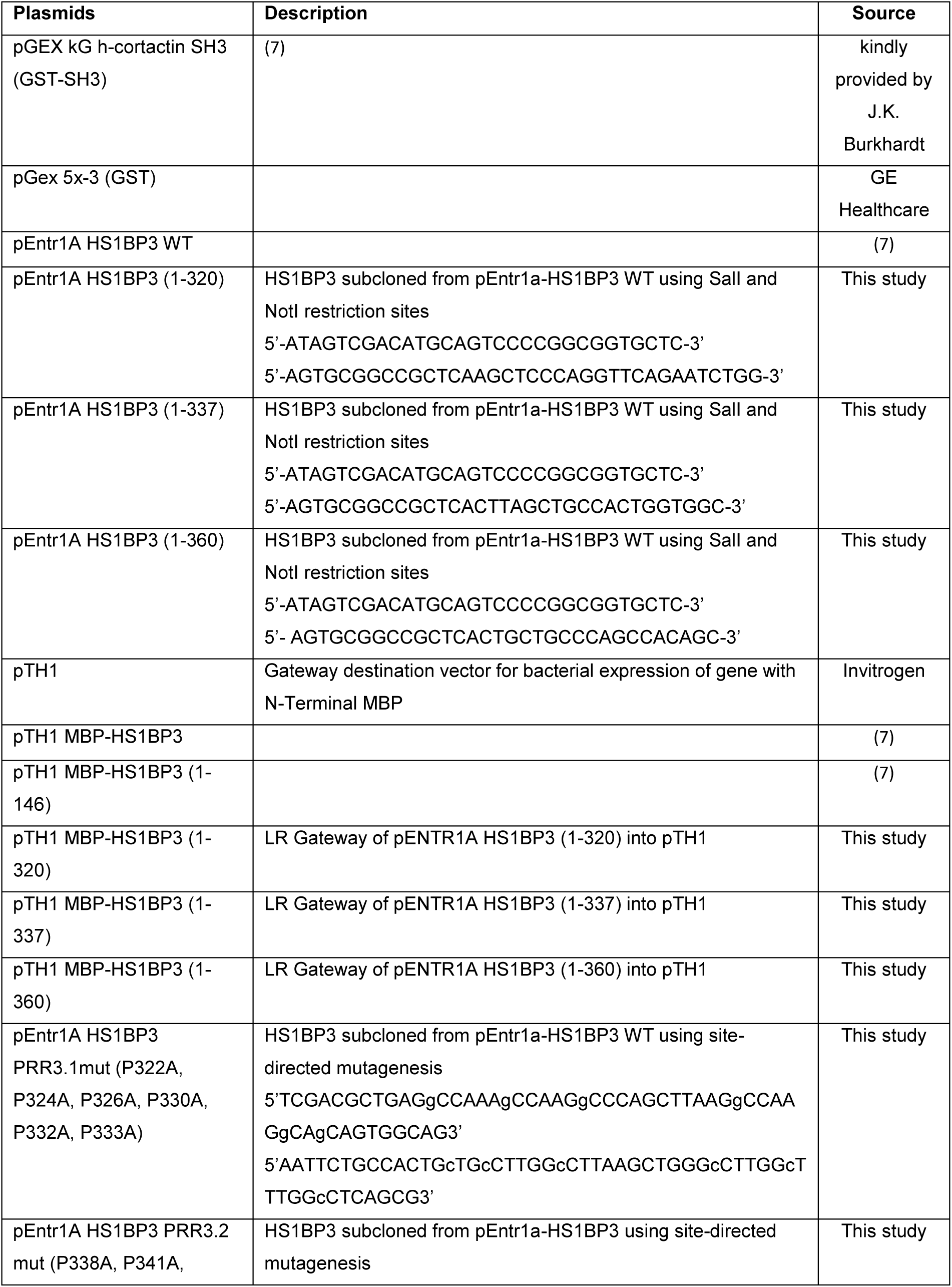
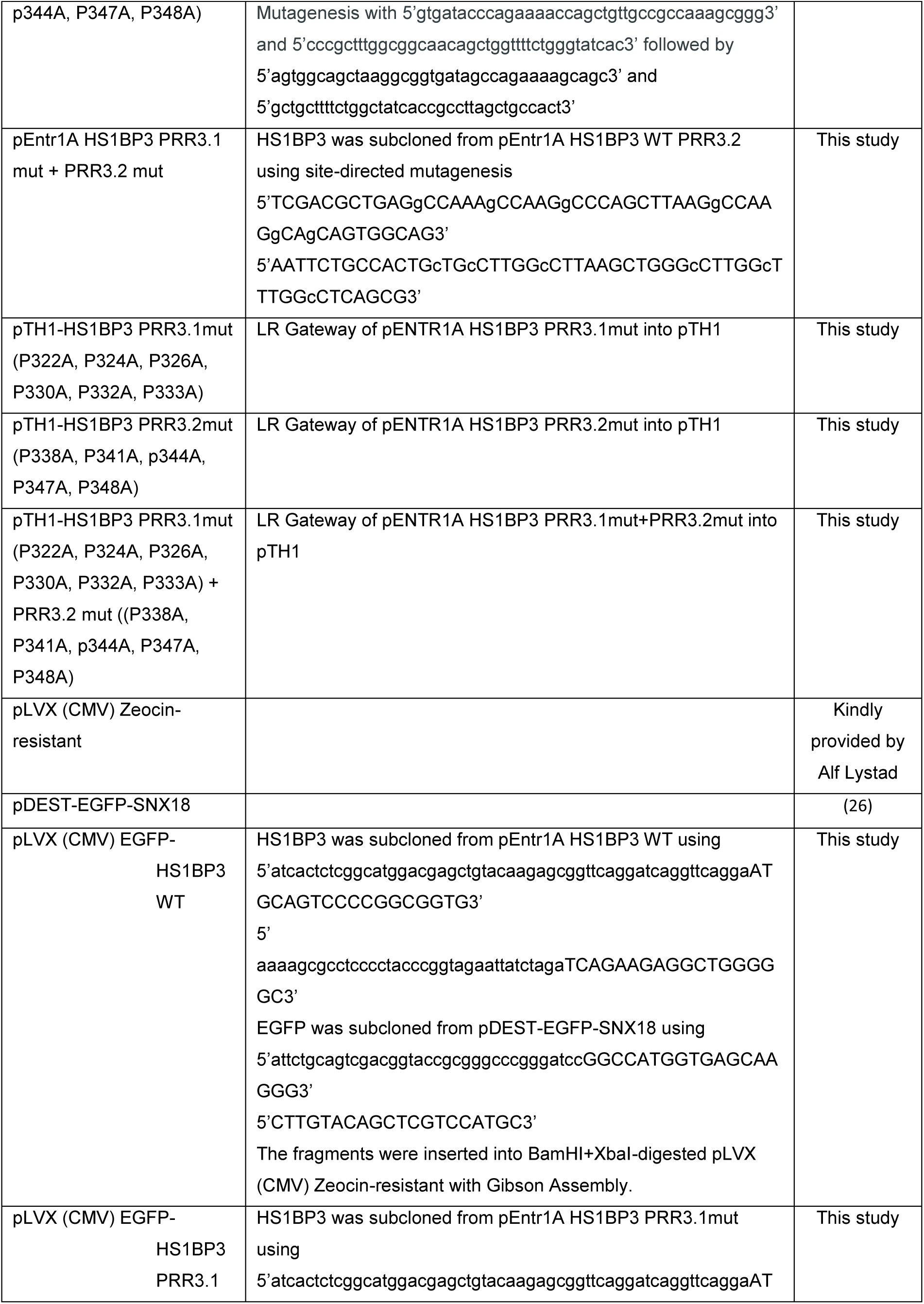
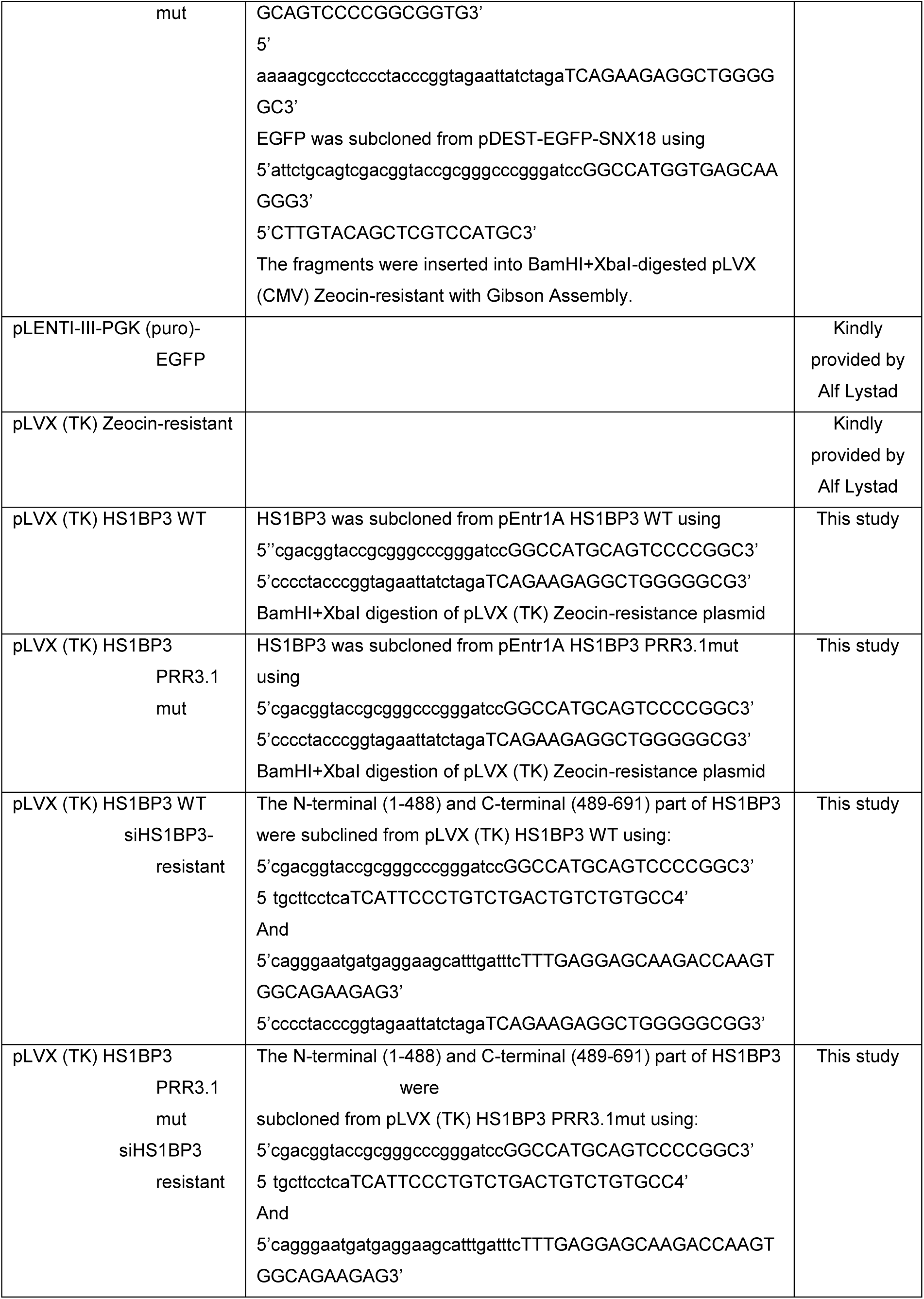
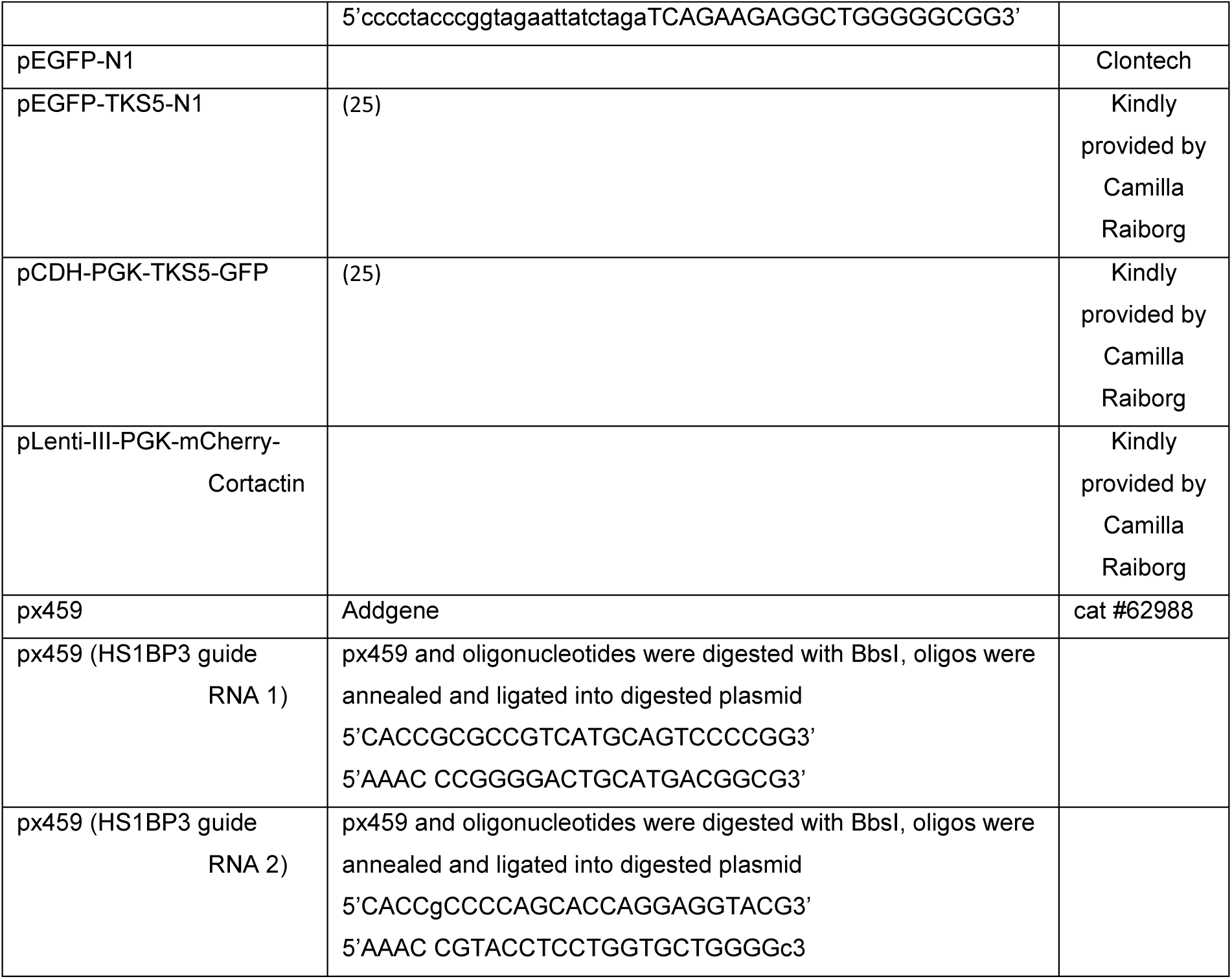
Overview of the oligonucleotides and cloning strategies used to generate the constructs used.

